# Capsular surveillance-guided phage cocktails against carbapenem-resistant *Klebsiella pneumoniae*

**DOI:** 10.64898/2026.11.13.766362

**Authors:** Chao Wang, Jingyi Nie, Min Wei, Xueru Tian, Yurou Sun, Yuan Zeng, Dawei Wei, Yuqin Song, Gang Zhang, Xiaobing Chu, Qiwen Yang, Zhemin Zhou, Linhuan Wu, Jie Feng

**Author notes:** Corresponding authors. E-mail addresses. These authors contributed equally: Chao Wang, Jingyi Nie, Xueru Tian, Min Wei.

## Abstract

Carbapenem-resistant *Klebsiella pneumoniae* (CRKP) is among the most urgent antimicrobial-resistance threats, yet its extraordinary capsular diversity has kept phage therapy locked in per-isolate matching — a process too slow and labor-intensive for routine use. Here we convert this specificity problem into a look-up problem. Interrogating 26,138 CRKP genomes from 96 countries, we show that capsular diversity collapses onto a tractable target list: 14 capsular locus (KL) types dominate contemporary circulation and jointly account for 67–77% of the CRKP burden in every WHO region, stably from 2010 to 2024 and after removal of sampling bias. We assembled a repository of 48 strictly lytic phages spanning all 14 types and organized it into capsule-defined modules: for eight KL types a single capsule-targeting phage sufficed to prevent resistant regrowth over 24 h, and for the remaining six a two-phage module pairs a capsule-targeting phage with a capsule-independent phage that recognizes the second receptor layer exposed upon capsule loss. We treated 28 clinical isolates, covering two distinct STs for each of the 14 KL types. Modules were assigned from the KL call alone, without isolate-level matching data. Growth was suppressed in every strain, and ST background had no significant effect in 12 of 14 types. These results establish capsule type, not lineage, as the operative unit of CRKP phage preparedness: a single rapid KL-typing step converts a diagnostic result into a therapeutic decision, providing a practical, lineage-agnostic route from surveillance to treatment.

---

Keywords: Carbapenem-resistant *Klebsiella pneumoniae*; phage therapy; capsular locus type; phage cocktail; sequence type

## Introduction

Antimicrobial resistance (AMR) poses a major threat to human health. A recent report estimated that in 2021, 4.71 million deaths were associated with AMR, with 1.14 million deaths directly attributable to it [1]. Meanwhile, the antibiotic development pipeline is shrinking as more pharmaceutical companies opt out of discovering and developing new antibiotics [2]. Therefore, the development of alternative antibacterial therapies has become imperative.

Accumulating clinical cases attest to the efficacy of phage therapy as a personalized treatment [3,4]. Yet behind each success lies a workflow that keeps the approach at the margins of routine care. Treating a single patient typically requires screening large environmental collections or curated libraries for phages that lyse the specific isolate. This process takes days to weeks [5,6] and is followed by whole-genome safety reviews and expansion of the selected phage to therapeutic titer. Such delays can determine outcomes for acutely ill patients. Therefore replacing labor-intensive per-isolate matching with a rapid look-up strategy based on capsule typing is key to making phage therapy scalable for routine use.

Carbapenem-resistant *K. pneumoniae* (CRKP) is a critical WHO priority pathogen and a leading cause of healthcare-associated infections [7]. In this species, the capsule determines phage specificity. It masks surface receptors and blocks phages adapted to other capsular types [8–11]. More than 160 genetically distinct KL types exist, but most phages target only one or two [8–10,12,13]. This renders a universal cocktail impractical. However, KL prevalence varies by region. KL64 and KL47 dominate in China and often carry *bla*_KPC-2_ and *bla*_NDM_ [14]. KL107-related types are common in the Americas and parts of Europe [15]. Across all regions, a few types cause most infections [16]. This concentration reduces an apparently intractable diversity problem to a finite target set. A similar approach worked for carbapenem-resistant *Acinetobacter baumannii*, where identifying the dominant ST2-KL3 lineage guided targeted phage isolation [17,18]. Genomic KL typing is now standard in surveillance pipelines [12,13], and rapid diagnostic tools work in clinical settings [23]. Yet current phage banks still require per-isolate susceptibility testing, and broad cocktails depend on predictable target populations [19–22]. Neither provides a standardized product. No repository currently links a KL type directly to a validated treatment module.

To fill this gap, we combined global capsular genomics with systematic phage characterization to build a KL-defined modular phage resource for CRKP. We analyzed 26,138 CRKP genomes from 96 countries and identified 14 globally prevalent KL types that account for most disease burden across all WHO regions. We then characterized 48 strictly lytic phages and assembled them into capsule-specific cocktail modules based on their receptor-recognition properties. For eight KL types, a single phage prevented resistant regrowth. For the remaining six, we paired a capsule-targeting phage with a capsule-independent phage that binds the second receptor layer exposed after capsule loss. We validated this approach using 28 clinical isolates, representing two distinct sequence types per KL type. Treatment was assigned solely from the KL genotype. The pre-assembled modules suppressed growth in all isolates, and efficacy remained consistent within KL types regardless of sequence type background. Capsular diversity can therefore be treated as a defined target list rather than an obstacle. This outlines a practical pathway from surveillance to treatment for CRKP without per-patient phage matching.

## Results

### A fourteen-type target list concentrates the CRKP burden in every WHO region

To establish an epidemiological framework of capsular type, we initially retrieved 71,132 *K. pneumoniae* genomes from the NCBI database (accessed through August 2024), spanning 122 countries across all six WHO regions. After stringent quality control (sequence completeness verification, metadata validation, and removal of redundant and sampling-biased genomes), we retained 45,050 high-quality genomes with complete spatiotemporal annotations. Resistance gene annotation identified 142 carbapenem resistance-associated genes, excluding both efflux pump-mediated resistance genes and outer membrane porin (OMP) genes. Isolates harboring at least one carbapenem resistance gene were designated CRKP, yielding a final dataset of 26,138 high-quality genomes from 96 countries. Carbapenemase-encoding genes dominated this profile, accounting for 92% (130/142) of identified determinants, and carbapenemase carriage was reported to strongly correlate with elevated meropenem MICs [25]. The proportion of CRKP among sequenced *K. pneumoniae* genomes varied considerably across WHO regions: the AMRO region had the highest proportion (66.4%) and the AFRO region the lowest (26.7%), while EMRO, SEARO, EURO, and WPRO accounted for 57.4%, 60.0%, 57.0%, and 54.7%, respectively (Figure S1A). We also quantified the contribution of each region to the global CRKP burden: AMRO accounted for the largest proportion of CRKP isolates (31.15%), followed by EURO (29.12%), WPRO (28.41%), SEARO (6.37%), EMRO (3.36%), and AFRO (1.59%) (Figure S1B). This geographic skew mirrors both regional differences in prevalence and uneven genomic sequencing intensity.

Kleborate predicted 127 distinct capsule synthesis loci (KL types) among these CRKP genomes. Applying an epidemiological criterion of ≥2% relative abundance in any WHO region identified 23 high-prevalence KL types (Figure 1A). Of these, 14 KL types (KL64, KL51, KL107, KL25, KL2, KL102, KL47, KL15, KL106, KL24, KL17, KL112, KL10, and KL62) exhibited global dissemination (≥2% prevalence in at least two regions) and were defined as globally prevalent. These 14 KL types accounted for 67.39–76.68% of the CRKP burden within each WHO region, indicating ecological fitness advantages that persist across geographical settings; KL64 dominated in all six regions, with prevalence ranging from 9.62% (AFRO) to 35.39% (WPRO). Temporal analysis (2010–2024) revealed that these types consistently constituted 59–79% of annual global CRKP isolates, persistently exceeding 50% and underscoring their sustained global dissemination (Figure S2). Recalculation before sampling-bias removal (based on Mash distance ≤0.00001) left the high prevalence of all 14 KL types unchanged, confirming that the target list is robust to surveillance structure.

**Figure 1.**
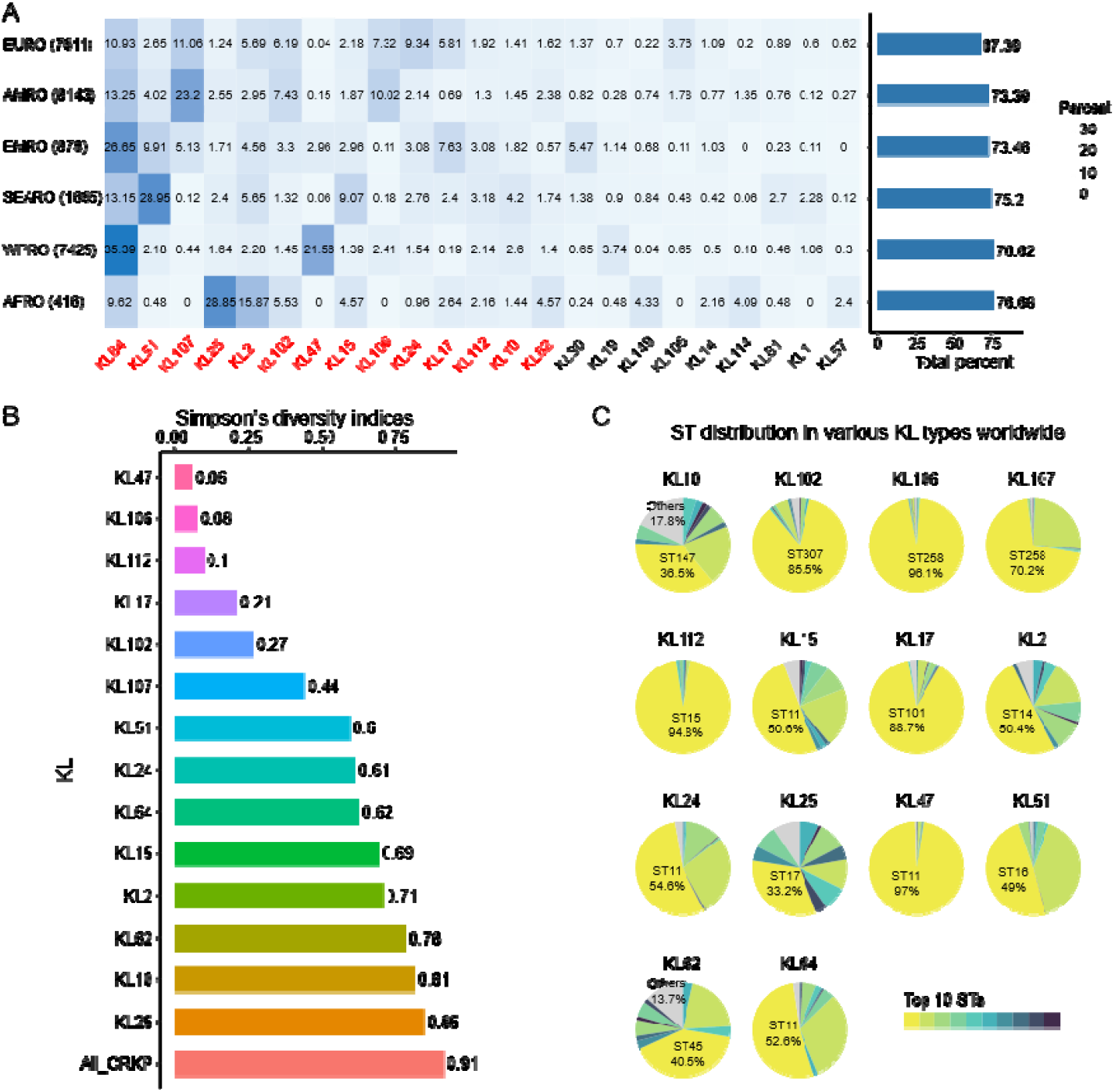
Global prevalence of KL types and ST diversity in CRKP. A. Regional distribution of high-prevalence KL types in CRKP. Left: heatmap of relative KL prevalence across the six WHO regions—EURO, AMRO, EMRO, SEARO, WPRO and AFRO; the genome counts for each region are indicated in parentheses after the region labels on the left vertical axis, and deeper color indicates higher prevalence. KL types classified as globally prevalent are highlighted in red. Right: bar chart summarizing, for each region, the cumulative prevalence of all globally prevalent KL types. B. Simpson’s diversity indices (SDIs) of ST distributions within the 14 globally prevalent KL types; the bar length represents the value of the SDI, with exact values indicated. C. ST composition of the 14 globally prevalent CRKP KL types. The sector size in each pie chart represents the proportion of genomes belonging to a given ST. The top 10 STs are shown in distinct colors, and the gray sectors represent all other STs, i.e., all STs in the 14 globally prevalent CRKP KL types other than the top 10. Abbreviations: CRKP, carbapenem-resistant *Klebsiella pneumoniae*; KL, capsular locus; ST, sequence type; SDI, Simpson’s diversity index; WHO, World Health Organization; EURO, European Region; AMRO, Region of the Americas; EMRO, Eastern Mediterranean Region; SEARO, South-East Asia Region; WPRO, Western Pacific Region; AFRO, African Region.

To characterize the multidrug resistance of the 14 globally prevalent KL-type clones, we profiled their antibiotic resistance genes (ARGs) content. Over the past two decades, the average number of ARGs per genome within these KL-type clones has remained relatively stable (31–35 ARGs/genome), slightly exceeding that of the overall CRKP population (30–34 ARGs/genome) (Figure S3). Most strains of these KL types harbor ARGs conferring resistance to β-lactams (>93%) and aminoglycosides (>77%), whereas colistin resistance genes are rarely detected (Table S1). By contrast, ARGs against macrolides, fluoroquinolones, tetracyclines, and chloramphenicols are heterogeneously distributed across KL types (17.49–78.28%, 2.96–65.11%, 5.41–65.82%, and 9.98–74.74%, respectively). KL107 and KL106, for instance, carry few tetracycline (<10%) and fluoroquinolone (<5%) resistance genes yet abundant chloramphenicol (>63%) and macrolide (>55%) resistance genes. KL62, similarly, carries ARGs against five antibiotic classes but few chloramphenicol resistance genes (13.5%). Thus, multidrug resistance is convergent across KL types, but its composition varies by type.

### Globally prevalent CRKP KL types exhibit divergent ST diversity and regional clonal patterns

We observed high diversity in the sequence types (STs) across KL types in CRKP, with an overall Simpson’s diversity index (SDI) of 0.91 and a total of 968 STs identified. The 14 globally prevalent CRKP KL types fell into 340 distinct STs, with SDIs ranging from 0.06 to 0.85. We identified 8 to 57 STs per type, reflecting substantial ST heterogeneity (Figure 1B). Most KL types were dominated by a single major ST, although the relative proportions varied. In KL47, ST11 accounted for 97.0% of isolates; in KL106, ST258 accounted for 96.1%; and in KL112, ST15 accounted for 94.8%. For the remaining KL types, the leading STs contributed between 36.5% and 88.7% of isolates. ST11 was the most prevalent ST within KL15, KL24, KL47, and KL64 (Figure 1C).

To compare the ST distribution of the same KL type across WHO regions, we calculated the number of STs and Simpson’s diversity index per region (Figure S4A, Table S2) and the similarity of ST distributions between regions (Figure S4B). ST distributions varied across WHO regions for most KL types: 11 of the 14 types showed high diversity (SDI >0.6) in at least one region (Figure S4, Table S2). There is strong regional heterogeneity in the ST components of KL107, KL47, KL25, and KL62, with significant differences in ST types and numbers between regions (similarity value <0.5 between any pair of regions) (Figure S4B). In contrast, no KL type showed high similarity of ST distribution across all regions. KL51, KL112, KL15, and KL17 showed relatively consistent ST distributions across regions (pairwise similarity >0.5 among three or four WHO regions). The remaining KL types exhibit transitional characteristics. Thus no single ST dominated all regions for any globally prevalent KL type. The chromosomal backgrounds beneath a given capsule are numerous and geographically fluid. This heterogeneity is precisely what a capsule-indexed product must tolerate to be deployable.

### A 48-member strictly lytic phage repository covers all fourteen target KL types

Using the previously generated host-range dataset for our *K. pneumoniae* phage library (178 strictly lytic isolates) [8,26], we identified 48 phages that reproducibly formed clear lysis zones on at least one member of the 14 globally prevalent KL types (Table S3). To quantify host range more rigorously, we assembled a panel of 65 representative *K. pneumoniae* isolates (2–7 isolates per KL type) (Table S4) covering the 14 target KLs. The resulting phage–host matrix comprised 3,120 possible pairwise interactions. Spot assays identified 366 (11.73%) lysis-positive pairs, of which 172 (5.51%) formed plaques with progeny production upon validation by double-layer agar plating (Figure 2A). Of the 65 tested strains, 58 were susceptible to at least one phage with confirmed progeny production. For the remaining 7 strains, clear lysis zones formed in spot assays, yet no progeny-producing phage could be recovered. Every KL type in the panel was susceptible to at least one phage. KL2 and KL64 supported the highest number of productive phages (6–14 and 2–10, respectively), whereas KL17, KL24, KL51, KL102, KL106, KL25, and KL107 were the most refractory. After removing non-productive lysis events, the matrix showed clear modularity.

**Figure 2.**
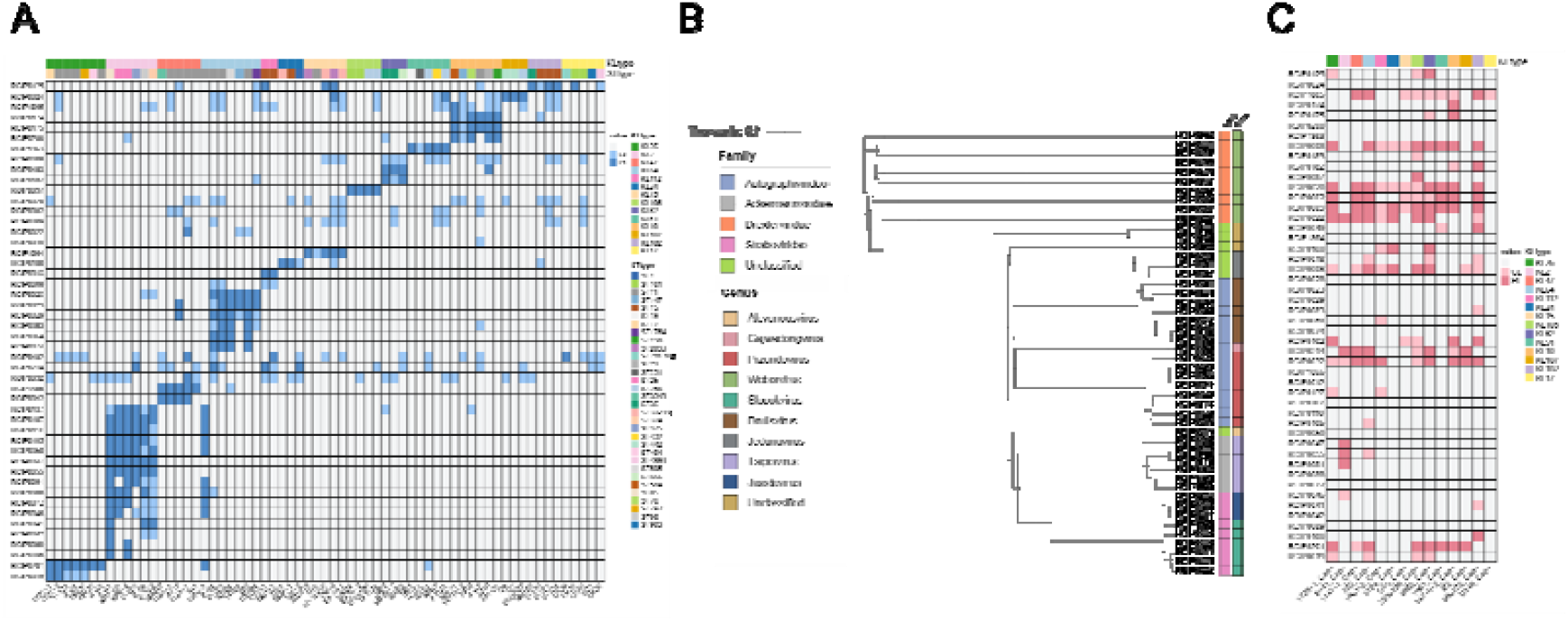
The phage–bacteria infection matrix. A. Phage–wild-type (WT) strain interaction matrix, in which 65 *K. pneumoniae* WT strains (columns) were exposed to 48 phages (rows). Blue squares denote spot-assay lysis; dark blue squares denote productive interactions (PI) confirmed by double-layer agar plating (plaque formation and progeny production); light blue squares denote clearing without confirmed productive infection (CL). Colored strips above the matrix categorize the *K. pneumoniae* strains by KL type. B. Neighbor-joining phylogenetic tree of the 48 phages, based on the Dice similarity coefficient of shared proteins, with color strips marking the taxonomic classification of the phages, with the left strip the family, and the right strip representing the genus. C. Interaction matrix of the 14 acapsular mutant (Cap) strains exposed to the 48 phages. Red squares indicate phage-mediated lysis of Cap strains evidenced by visible plaque formation; dark red squares denote productive interactions (PI); light red squares denote clearing without confirmed productive infection (CL). Colored strips above the matrix categorize the strains by KL type.

Whole-genome sequencing showed that the 48 lytic phages are taxonomically diverse, spanning four families and nine recognized genera within the class *Caudoviricetes* (Figure 2B). *Autographviridae* dominated the collection (16/48 phages), distributed across three genera — *Drulisvirus* (n = 7), *Przondovirus* (n = 8) and a single *Gajwadongvirus*. Ten phages grouped within *Drexlerviridae*, all assigned to the genus *Webervirus.* Nineteen phages were distributed across further taxa: three clustered into a distinct *Jedunavirus* clade, nine were assigned to the family *Straboviridae* (three *Jiaodavirus* and six *Slopekvirus*), six belonged to the family *Ackermannviridae* (genus *Taipeivirus*), and one could only be categorized at the genus level as *Alcyoneusvirus*. In addition, three phages remain unclassified beyond the class level. Notably, none of the genomes encoded integrases, repressors, toxins, or antibiotic resistance determinants (Table S3), confirming a strictly lytic lifestyle and reinforcing their therapeutic potential [27]. The breadth of capsule tropism did not align with phylogenetic relationships: phages within the same genus often targeted different KL types, whereas distantly related phages occasionally converged on the same capsule type, suggesting that KL specificity is driven by receptor-binding proteins rather than overall genomic lineage.

To identify the molecular determinants of the KL-type-specific lysis spectrum, we focused on tail fiber proteins predicted to act as depolymerases — receptor-binding proteins that specifically recognize and degrade capsular polysaccharides [28]. Depolymerase-carrying tail fibers were identified in 32 of the 48 phages, spanning six genera: *Drulisvirus* (n = 7), *Przondovirus* (n = 7), *Webervirus* (n = 8), *Jedunavirus* (n = 3), *Alcyoneusvirus* (n = 1) and *Taipeivirus* (n = 6). We modeled each predicted depolymerase as a monomer with AlphaFold3 [29] (Figure S5). All 32 models exhibited the canonical architecture of an N-terminal anchoring domain and a central β-helical catalytic domain, confirming their capacity for capsular polysaccharide recognition and degradation.

### Precision phage cocktails tailored to globally prevalent CRKP KL types

Therapeutic phages are usually administered as cocktails — defined, complementary mixtures that slow or prevent the emergence of resistant mutants [19,20,26]. Because the CPS forms the outermost barrier and serves as the primary receptor for most *K. pneumoniae* phages, we deliberately paired CPS-dependent with CPS-independent phages for every clinically relevant KL type. To generate receptor-diversified targets, fourteen representative clinical isolates were serially passaged; spontaneous acapsular (Cap) variants emerged for all fourteen isolates (Figure S6; Table S5). We next screened the 48 characterized lytic phages against all Cap derivatives. Thirty-one phages readily lysed these acapsular mutants, providing at least one CPS-dependent and one CPS-independent candidate for each of the 14 evaluable KL types (Figure 2C). Based on this host-range matrix, we formulated receptor-diversified modules for each isolate and quantified their antibacterial activity in 24-h turbidimetric assays at an MOI of 1.0.

We first characterized the *in vitro* lysis kinetics of the 14 KL-type strains, each treated with the capsule-targeting (CPS-dependent) phage(s) matched to its own KL type. One KL-matched phage per type was sufficient for eight KL types (KL2, KL17, KL24, KL25, KL102, KL106, KL112, and KL47): bacterial density remained low with no resistant regrowth over 24 h (Figure 3A–H). For KL24, KL25, and KL106, the effective phage also lysed acapsular mutants of the same strains, indicating that it carries multiple receptor-binding proteins (RBPs). For KL2, KL17, KL102, KL112, and KL47, the KL-matched CPS-dependent phage alone held bacterial density near the starting level at 24 h post-infection.

**Figure 3.**
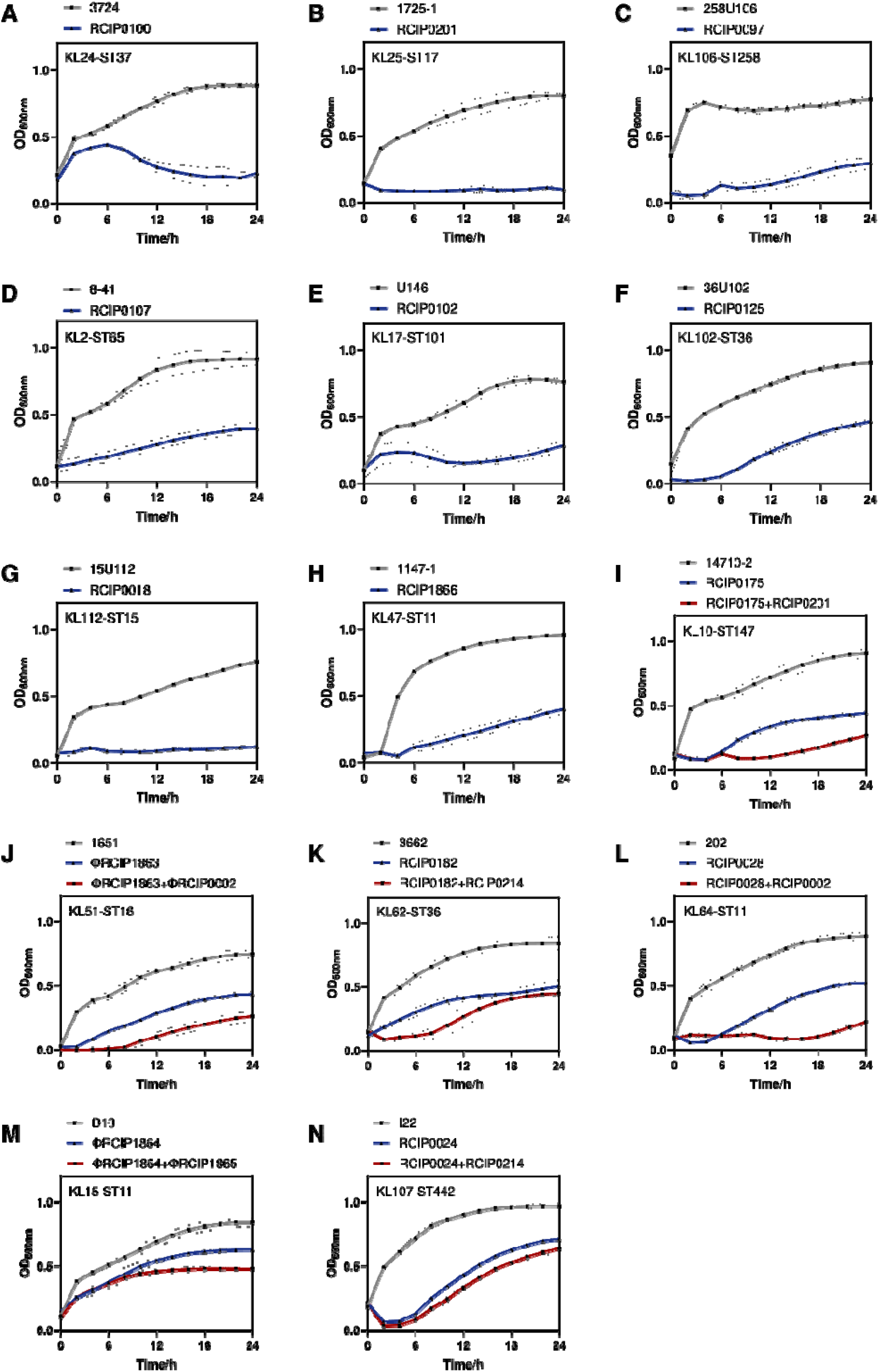
Growth curves for untreated controls and 14 *K. pneumoniae* KL-type isolates treated with phages. Each panel shows OD_600_ over 24 h for the untreated control and the phage cocktail at an MOI of 1.0. (A–H) Present the eight KL types treated with a single capsule-dependent phage (KL2, KL17, KL24, KL25, KL102, KL106, KL112, and KL47). (I–N) Present the six KL types treated with dual-phage cocktails (KL10, KL51, KL62, KL64, KL15, and KL107). Data are means ± SD of three independent replicates.

The remaining six KL types developed resistance to their capsule-targeting phages (Figure 3I-N). We therefore paired each KL-matched capsule-targeting phage with a non-capsule-targeting phage. For KL10, KL51, KL62, and KL64, the dual-phage modules suppressed bacterial growth throughout the 24 h and delayed regrowth by at least 12 h compared with the corresponding monophage treatment. For KL15, neither the capsule-targeting nor the non-capsule-targeting phage produced progeny on this strain, but the two-phage combination still suppressed bacterial growth. For KL107, the dual-phage module showed no significant advantage over the single phage, yet optical density (OD_600_) remained below 0.5 at 24 h, in contrast to the phage-free control group, which grew to high density.

### Deployment by KL look-up with phage assigned from the capsule type alone suppresses growth across 28 isolates of diverse sequence types

The repository is only useful as a product if treatment can be assigned without re-running phage–host matching for each new isolate. We therefore tested the look-up rule directly. Each of the 14 KL types received its pre-defined KL-matched treatment — a single capsule-dependent phage for eight types, and a two-phage capsule/capsule-independent module for the remaining six (KL64, KL51, KL107, KL15, KL10, and KL62) — and each treatment was applied to two clinical isolates of distinct STs within its KL type (28 strains in total), in 24 h turbidimetric assays at an MOI of 1.0. The treatment choice for every strain was determined solely by its KL type; isolate-level host-range data were not used to alter module assignment. Efficacy was quantified as ΔAUC, the difference in area under the growth curve relative to the untreated control. In all 28 strains, the KL-matched treatment suppressed growth significantly below the untreated control (Figure 4A).

**Figure 4.**
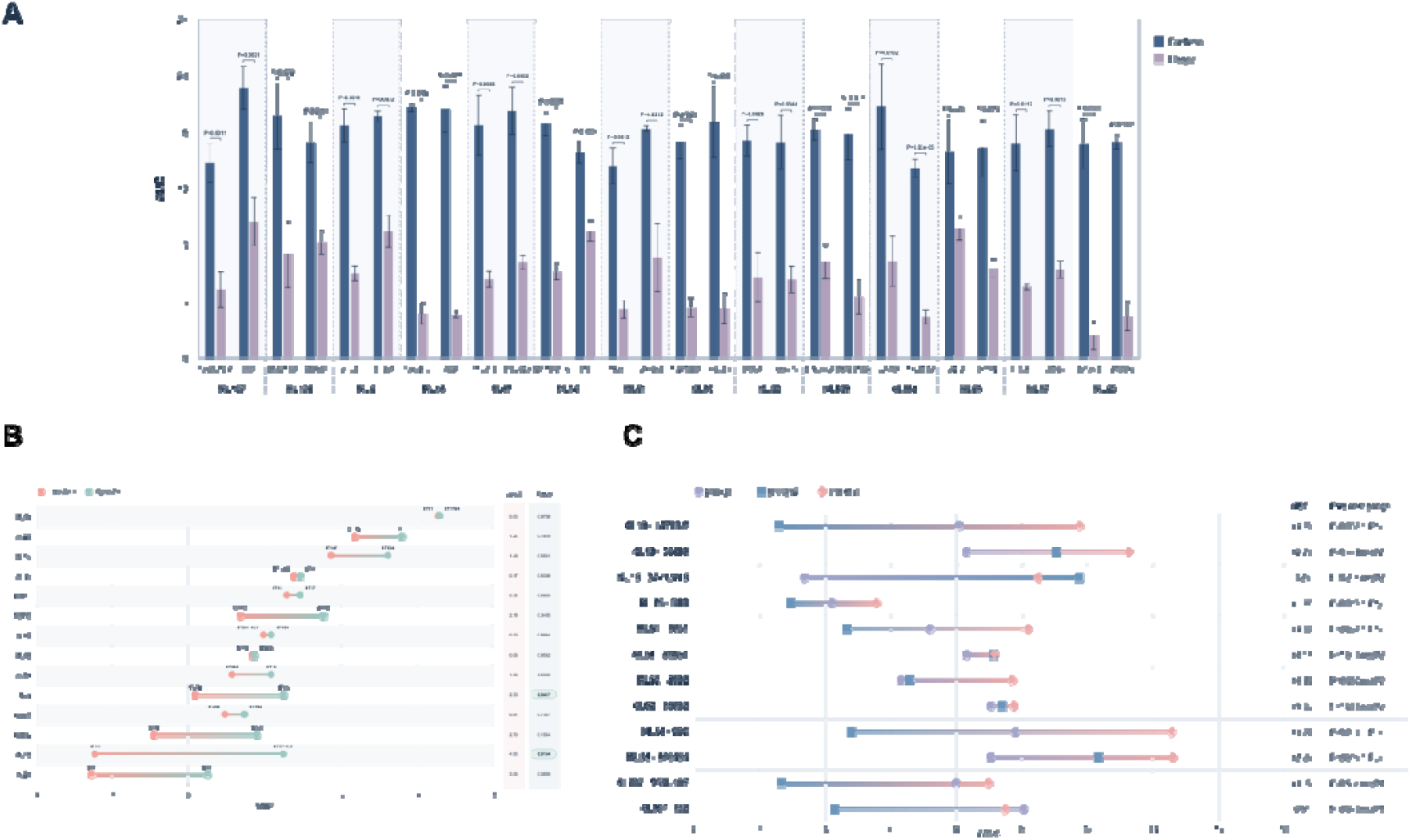
Efficacy of KL-matched treatments against CRKP across KL types and STs, and comparison of dual-phage cocktails with their constituent phages. (A) Growth-inhibitory efficacy of the KL-matched treatment against 28 CRKP strains (two strains of distinct STs per KL type; 14 KL types), measured in 24-h turbidimetric assays at an MOI of 1.0. The treatment comprised a single capsule-dependent phage for eight KL types (KL2, KL17, KL24, KL25, KL102, KL106, KL112 and KL47) and a two-phage cocktail pairing a capsule-dependent with a capsule-independent phage for the remaining six (KL64, KL51, KL107, KL15, KL10, and KL62). ΔAUC denotes the difference in the area under the OD600 growth curve (AUC) relative to the untreated control, such that larger values indicate stronger growth inhibition. Bars show means of three independent replicates, with P values (Welch two-sided t test, treatment versus untreated control) shown above the bars. B. Influence of ST background on treatment efficacy within each KL type. For each of the 14 KL types, the ΔAUC values of the two strains of distinct STs are compared. The between-ST difference in treatment efficacy is expressed as ΔΔAUC = ΔAUC(strain A) − ΔAUC(strain B), where strain A and strain B are the two strains of the same KL type, and its absolute value |ΔΔAUC|, together with the P value of the between-ST comparison, is indicated. |ΔΔAUC| quantifies the magnitude of the between-ST difference in efficacy regardless of direction; larger values indicate that treatment efficacy depends more strongly on the ST background. Between-ST differences reaching the nominal threshold (P < 0.05, highlighted) were restricted to KL2 and KL15; the remaining 12 KL types showed no significant difference. C. Component contributions of the dual-phage cocktails. ΔAUC of the capsule-dependent phage (phage 1), the capsule-independent phage (phage 2), and the two-phage cocktail (phage 1+2) for the 12 strains of the six dual-cocktail KL types (two strains per KL type), together with the gain of the cocktail over the better single phage, quantified by the highest-single-agent (HSA) model: HSA = ΔAUC(cocktail) − max[ΔAUC(phage 1), ΔAUC(phage 2)]. HSA > 0 indicates that the cocktail inhibited growth more strongly than the better-performing single phage. Within each strain, the cocktail was compared with its better-performing constituent phage by two-sided Welch’s t tests; the P values are nominal (unadjusted for multiple comparisons), and cocktail performance was assessed primarily by the HSA effect size. All data are means of three independent replicates. Two-sided P < 0.05 was considered statistically significant; ns, not significant. Statistical analyses were performed in R. Abbreviations: CRKP, carbapenem-resistant *Klebsiella pneumoniae*; KL, capsular K-locus type; MOI, multiplicity of infection; ST, sequence type.

To test whether the ST background of a strain alters the outcome of a KL-assigned treatment, we compared ΔAUC between the two STs within each KL type. Of the 14 KL types, 12 showed no significant between-ST difference (Figure 4B). The two exceptions were KL2 and KL15: within KL2, strain 8-41 (ST65) had a higher mean ΔAUC than strain 8-152 (ST374) (10.49 vs. 8.16; absolute difference, 2.33; P = 0.0417), and within KL15, strain 37-1LV15 had a higher mean ΔAUC than strain D10 (ST11) (10.49 vs. 5.54; absolute difference, 4.95; P = 0.0154). KL24 and KL102 exhibited the largest non-significant differences (3.06, P = 0.0889; and 2.70, P = 0.1564 respectively). These findings indicate that KL-assigned treatment produces comparable growth inhibition across isolates with different ST backgrounds; strain- or ST-associated genetic factors may modulate the magnitude of response within certain KL types, but did not overturn the assignment rule in any case.

We next asked what the second component of each dual module contributes. Each two-phage module was compared with its two constituent phages in the 12 isolates representing the six dual-module KL types (Figure 4C). The module significantly outperformed the better single phage in five isolates: KL10 strain 14710-2 (HSA = 3.69, P = 0.028), KL15 strain D10 (HSA = 1.35, P = 0.012), KL51 strain 1651 (HSA = 3.00, P = 0.037), KL64 strain 202 (HSA = 4.78, P = 0.011), and KL64 strain 176464 (HSA = 2.29, P = 0.019). The two KL64 isolates, from distinct STs, both benefited significantly. KL62 strain 3662 showed a large but non-significant gain (HSA = 3.13, P = 0.060). In two isolates the module was numerically less effective than the better single phage, though not significantly: KL15 strain 37-1LV15 (HSA = −1.25) and KL107 strain I22 (HSA = −0.57). The two-phage module matched or exceeded the better single phage in 10 of the 12 isolates, with significant gains in five and no significant loss in any: adding the capsule-independent component preserved, and in some strains improved, efficacy while removing the escape route of a single receptor layer.

## Discussion

Our findings demonstrate that the extraordinary capsular diversity of CRKP can be effectively managed through a capsule-defined modular framework. The identification of 14 globally prevalent KL types that account for the majority of the burden across all WHO regions highlights a critical epidemiological feature: stability amidst heterogeneity. Although regional rankings of these types vary, the core set remains consistent over fifteen years. This suggests that phage preparedness should focus on this finite target list rather than attempting to cover the entire spectrum of 160+ KL variants. Such an approach mirrors successful strategies in other pathogens, such as *Acinetobacter baumannii*, where targeting dominant lineages enabled scalable intervention [17,18]. By converting the specificity problem into a look-up task, we provide a rationale for prioritizing surveillance resources toward these high-impact KL types.

The design of our cocktail modules accounts for resistance evolution, not just initial susceptibility. For eight KL types, no resistant regrowth was observed within the 24-hour window, indicating that single-phage modules are sufficient to suppress population recovery. For the remaining types, we employed a two-phage strategy pairing a capsule-targeting phage with a capsule-independent one. This addresses escape via capsule loss. Although acapsular strains suffer from attenuated virulence in vivo [30,31], they can still emerge under selective pressure. Pairing phages that target distinct surface layers exploits cross-resistance trade-offs, a strategy we previously described for overcoming phage resistance[26]. By forcing bacteria to accumulate multiple independent genetic changes, this approach significantly raises the barrier to resistance and explains why our modules maintained efficacy without frequent emergence of fully resistant clones.

A pivotal insight from this study is the dissociation between chromosomal background and therapeutic outcome. We observed substantial ST polymorphism within each KL type, yet treatment efficacy remained conserved regardless of sequence type. In 12 of the 14 KL types, differences in growth suppression between distinct STs were statistically insignificant. This challenges the traditional view that strain-level matching is necessary for precision therapy. Instead, it establishes capsule type as the operative unit of phage preparedness. The selective advantage conferred by specific capsular structures appears to outweigh the influence of the underlying genomic lineage. Consequently, clinicians do not need to track regionally variable ST distributions to select a treatment. Combined with rapid KL-typing methods that enable same-day capsule identification [23], this lineage-agnostic property makes it feasible to retrieve the appropriate validated module directly from a diagnostic result.

Despite these promising results, several limitations warrant consideration. First, our repository covers the most prevalent KL types but leaves a residual gap for the 23–33% of infections involving less common variants; broader coverage will require continuous surveillance-driven updates. Second, although three biological replicates per strain ensured reproducibility, the limited number of distinct isolates tested for each KL type constrains our ability to detect subtle ST-specific effects, necessitating larger isolate panels before widespread clinical implementation. Third, receptor assignments currently rely on comparative genomics, and direct biochemical validation of second-layer receptor recognition is ongoing to confirm the mechanistic basis of our modules.

Looking forward, this framework offers a roadmap for translating phage therapy from bespoke experiments to standardized products. Continuous monitoring of KL prevalence ensures that the repository remains aligned with circulating strains, while periodic re-validation maintains product quality. Ultimately, defining the capsule as the key determinant enables scalable and precise phage therapy for CRKP, providing a template for pairing molecular epidemiology with phage-biology toolkits against other multidrug-resistant pathogens.

## Methods

### Bacterial isolates and growth conditions

*K. pneumoniae* isolates were grown at 37°C in Luria Bertani (LB) medium (10 g/L tryptone, 5 g/L yeast extract, and 10 g/L NaCl).

### Data collection and quality control

We collected 71,132 *K. pneumoniae* genomes as of August 2024 from the National Center for Biotechnology Information (NCBI) database. Completeness and contamination of genomes were checked using CheckM v1.2.2 [35]. Whole-genome Average Nucleotide Identity (ANI) with the reference genome was calculated using FastANI v1.33 [36]. Fast distance evaluation between any pair of genomes was performed using Mash v2.3 [37], followed by the mash sketch and mash dist commands. A self-written script was then used to cluster the genomes according to a distance ≤0.00001, with the representative genome in each cluster being the one with the smallest maximum distance from the other genomes within the cluster. Redundancy removal and clustering therefore preceded all prevalence estimates reported in the main text; a sensitivity analysis performed before redundancy removal gave the same prevalence patterns. Data were screened according to the following criteria: (1) completeness ≥95% and contamination ≤5% [38]; (2) ANI ≥95% [39]; (3) metadata on country of origin and time of isolation present; (4) contig number ≤1,000; (5) N50 ≥10,000; (6) removal of GC content outliers; (7) no genomes with duplicate biosample numbers; (8) KL type identification result “Typeable”; (9) genome annotated as “*Klebsiella pneumoniae*”; (10) representative genome of each cluster.

### KL type identification and gene annotation

Kleborate v3.0.9 [12] was used for KL typing, ST typing, and species prediction with the -p kpsc option. Antibiotic resistance genes (ARGs) were annotated with the Resistance Gene Identifier (RGI) v6.0.2 [40] rgi main with -t protein -a DIAMOND. Annotation results with identity ≥90% were selected. Protein sequences in the genome were identified using Prokka v1.14.6 [41] with parameters --gcode 11.

### Defining high-prevalence and globally prevalent KL types

According to the WHO, countries are classified into six world regions. Prevalence of each KL was calculated per region as its percentage among all KL calls in that region. A KL type was considered “high-prevalence” if its prevalence was ≥2% in at least one region, and “globally prevalent” if it met the same threshold in at least two regions. These thresholds and labels were defined independently of the concurrent study by Lei *et al.* [24], which uses continents as its geographic unit. We used WHO regions because they coincide with the administrative frame of the WHO bacterial priority pathogens list and of national surveillance reporting.

### Identifying CRKP

We extracted a list of carbapenem-related resistance genes from the Comprehensive Antibiotic Resistance Database (CARD) [40]. Because non-carbapenemase factors, including antibiotic efflux pumps and membrane porins, contribute relatively little to overall pathogenicity and clinical outcomes, genes associated with antibiotic efflux pumps and *K. pneumoniae* outer membrane porins were excluded from the definition of CRKP [14,42–44]. An isolate was considered CRKP if it carries at least one annotated resistance gene in this list.

### Resistance rates to various antibiotics

The proportion of CRKP genomes resistant to various antibiotics was calculated as the percentage of genomes harboring corresponding resistance genes relative to the total number of genomes, based on annotated ARG–antibiotic links in CARD. ARGs beginning with “CMY”, “CTX-M”, “OXA”, “SHV”, “TEM”, and “VIM” were classified as β-lactam resistance; ARGs beginning with “KPC”, “NDM”, “GES”, and “IMP” were excluded as universal in CRKP; “MCR” as colistin; “TET” as tetracycline; “Qnr” as fluoroquinolone; “AAC”, “aad”, “ANT”, “APH” as aminoglycoside; “cat” or “cmlA” as chloramphenicol; “Erm” and “mph” as macrolide resistance. All classifications were case-insensitive.

### Host-range determination and validation of lytic activity

The host range of the 48 isolated phages was evaluated against a panel of 65 wild-type *K. pneumoniae* clinical isolates representing the selected KL types (2–7 isolates per KL type; Table S4). Initial screening used a qualitative spot assay: 3 μL of high-titer phage lysate (∼10^9^ PFU/mL) was spotted onto a bacterial lawn followed by co-incubation on LB agar for 18 h at 37°C; spots exhibiting clear zones of lysis were recorded as positive. To distinguish true productive infection from artifacts such as lysis from without, all spot-positive interactions were validated by the double-agar overlay plaque assay [45]. Serial dilutions of phage preparations were mixed with the respective host bacteria in molten soft agar and overlaid onto solid LB agar. A phage–host combination was considered confirmed only if discrete plaques were reproducibly observed across serial dilutions. Lytic spectra were visualized as heatmap incidence matrices using the Heatmap function in the ComplexHeatmap (version 2.26.1) R package [46].

### Generation and genomic analysis of capsule-deficient mutants

To generate CPS-deficient mutants, isolates of 14 different KL types were serially passaged in liquid LB medium. The same 48 phages were tested against the 14 acapsular (Cap) mutants by spot assay, and all spot-positive interactions were verified by double-layer agar plating. Genomic DNA of all Cap mutants and their corresponding wild-type strains was extracted with a TIANGEN genomic DNA purification kit according to the manufacturer’s instructions and whole-genome sequencing was performed on the PromethION platform (NextOmics) and the NovaSeq 6000 PE150 platform (Novogene) to generate Nanopore long reads and Illumina paired-end short reads, respectively. Complete genomes were assembled *de novo* from Nanopore long reads using Flye and polished with Illumina paired-end reads using NextPolish. For the capsulated parental strain and its capsule-deficient phage-resistant mutants, Kaptive v3.0.0b5 [13] was run with the assembly command to delineate the capsular polysaccharide synthesis locus (K locus, KL) and identify KL-associated genes and their genomic coordinates. Other genes and their positions were annotated with Prokka v1.13.1 [41]. Single-nucleotide variants (SNVs) and indels were identified by aligning capsule-associated loci and whole-genome sequences against the parental genome with nucmer, show-snps -l -q and show-coords -rcl in MUMmer v4.0.0rc1 [47], alongside BLASTN from BLAST+ v2.16.0 [48] (-task blastn).

### Phylogenetic analysis of phages

Based on the protein sequences of the 48 Klebsiella phages, an all-versus-all comparison was performed with DIAMOND [50] (v2.0.7) in more-sensitive mode. A Dice distance matrix was then computed as previously reported [51], and used to construct a neighbor-joining tree in the “ape” R package (version 5.5) [52].

### Receptor-binding protein (RBP) and depolymerase analysis

RBPs were predicted and functionally annotated for all 48 phage genomes using homology-based searches [8,23]. Depolymerase genes were identified by homology to known depolymerase domains (e.g., pectate lyase, glycoside hydrolase and tail spike domains), and phages targeting the same KL type were compared for shared depolymerase homologues. Three-dimensional models of RBPs were predicted using AlphaFold 3 [29], with interface confidence evaluated by the predicted template modeling (ipTM) score.

### Bacterial growth curves under phage pressure

To assess *in vitro* lytic efficacy, an automated growth curve analyzer (Cerillo, Charlottesville, Virginia, USA) monitored bacterial cultures. In this assay, 100 μL of bacterial culture at OD_600_ = 0.2 (approximately 10^7^ CFU) was transferred to 100-well flat plates and mixed with phage solution: 100 μL of a single phage; 50 μL of each phage for two-phage modules; 33.3 μL of each for three-phage combinations; 25 μL of each for four-phage combinations, to a final MOI of 1.0 for each component. Controls were mixed with SM buffer (Coolaber, Beijing, China) without phage. OD_600_ was measured hourly for 24 h with periodic shaking. All experiments were performed in triplicate, and inhibition curves were plotted from mean OD_600_ values.

### Statistical analysis of growth curves

For growth-curve experiments, the area under the OD_600_ curve over 24 h (AUC) was computed per replicate using the trapezoidal rule [53]. Phage-mediated growth inhibition was quantified as ΔAUC = AUC(control) − AUC(treatment) [53]; larger positive ΔAUC indicates stronger inhibition. We report ΔAUC rather than the percentage-based Suppression Index of Kim *et al.* [53]; the two express the same AUC-based contrast in different units. To test whether phage susceptibility is conserved within capsular types, ΔAUC values were compared between isolates sharing a KL type but belonging to different STs. The between-ST difference per KL type was quantified as ΔΔAUC = ΔAUC(strain A) − ΔAUC(strain B) and reported as its absolute value in Figure 4B; a larger |ΔΔAUC| indicates stronger dependence of treatment efficacy on ST background. For each isolate, the two-phage module was compared with each constituent phage using the highest-single-agent (HSA, Highest Single Agent) model: HSA = ΔAUC(cocktail) − max[ΔAUC(phage1), ΔAUC(phage2)] [54]; HSA > 0 indicates the module inhibited growth more strongly than the better-performing single phage. Significance was assessed by comparing replicate ΔAUC values of the module against those of the better constituent phage.

### Statistical calculation

Simpson’s diversity indices were calculated with the diversity function of the vegan R package (index = “Simpson”). ST-distribution similarity was computed as 1 − Bray–Curtis dissimilarity using vegdist (vegan version 2.7-1) [55], and visualized with ggplot2 (version 3.4.2) [56]. For growth-curve data, comparisons between groups were performed using two-sided unpaired Welch’s t-tests. All reported P values are nominal and were not adjusted for multiple comparisons. In Figure 4B, P < 0.05 was used as the nominal threshold for highlighting between-ST differences. Cocktail performance in Figure 4C was assessed primarily by the HSA effect size, with Welch P values reported for descriptive comparison. All analyses were performed in R (version 4.5.0).

## Acknowledgements

We thank Dr Hui Wang, Department of Clinical Laboratory, Peking University People’s Hospital, Beijing, China, and Dr Jing Yuan, Children’s Hospital Capital Institute of Pediatrics, Chinese Academy of Medical Sciences & Peking Union Medical College, Beijing, China, for providing the microbial resource. This work received support from the National Key R&D Program of China (Grant No. 2025YFC3408400).

## Declaration of Competing Interest

The authors declare no conflict of interest.

## Ethics approval

This study used anonymized bacterial isolates obtained from existing collections. No human participants or human specimens were involved, and no patient-identifiable information was accessed; therefore, ethics approval was not required.

## Data availability statement

Genomic sequence data of the 48 phage genomes and the 65 *K. pneumoniae* isolates used in this study have been deposited in the National Microbiology Data Center (NMDC, https://nmdc.cn/). Public release is scheduled for 2027-10-01. The complete accession lists are provided in Tables S3 and S4.

## CRediT authorship contribution statement

Chao Wang: Writing – review & editing, Writing – original draft, Visualization, Validation, Software, Methodology, Formal analysis, Data curation, Conceptualization. Jingyi Nie: Writing – original draft, Methodology, Formal analysis, Data curation. Min Wei: Writing – original draft, Investigation, Methodology, Validation. Xueru Tian: Writing – original draft, Investigation, Methodology. Yurou Sun: Validation. Yuan Zeng: Visualization, Resources, Data curation. Dawei Wei: Supervision, Formal analysis. Yuqin Song: Software, Methodology, Data curation. Gang Zhang: Project administration, Supervision. Xiaobing Chu: Resources. Qiwen Yang: Resources. Zhemin Zhou: Writing – review & editing, Validation, Supervision. Linhuan Wu: Writing – review & editing, Validation, Supervision, Data curation, Conceptualization. Jie Feng: Writing – review & editing, Validation, Supervision, Project administration, Funding acquisition, Conceptualization.

---

**Figure S1.**
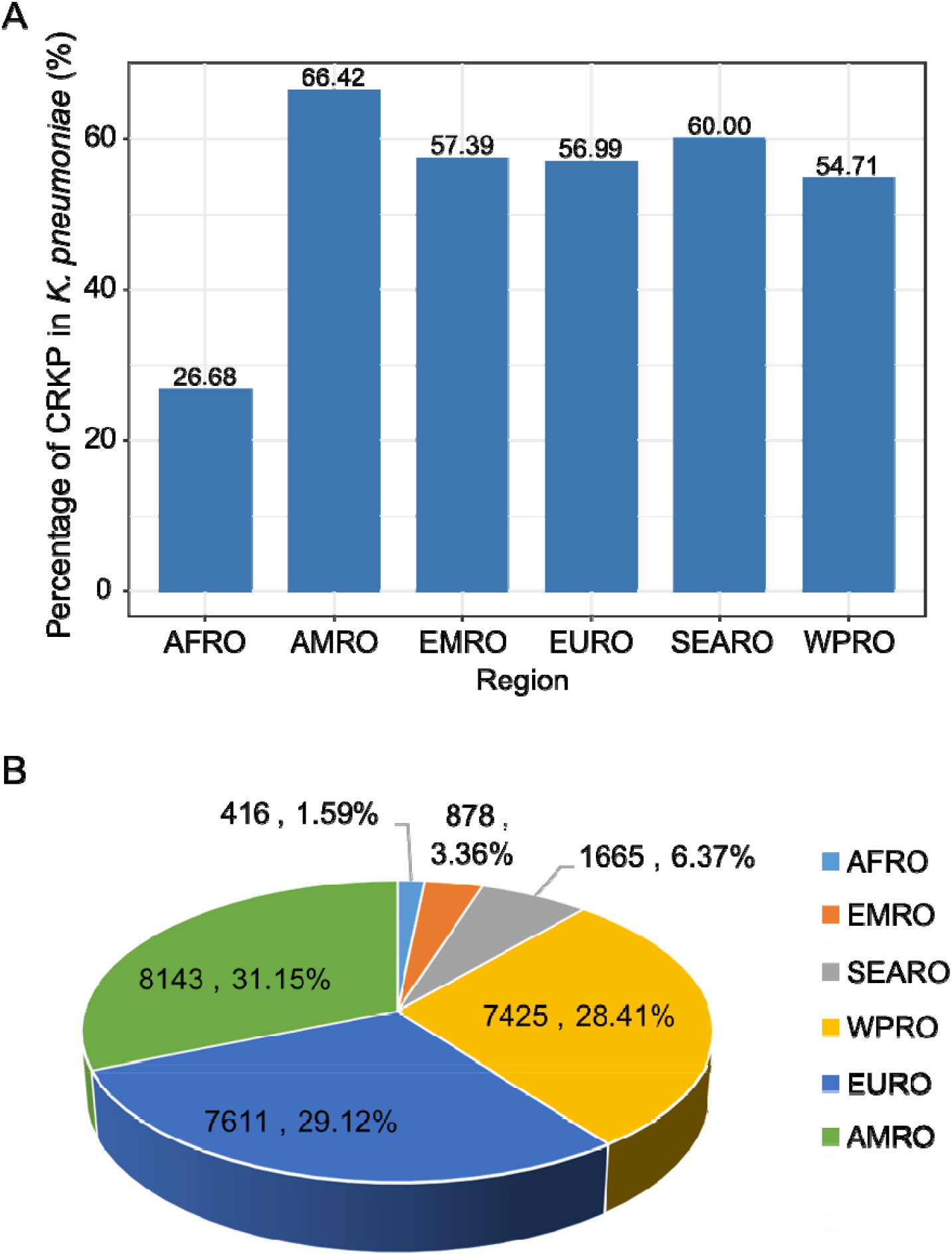
Regional distribution of carbapenem-resistant *K. pneumoniae* (CRKP). A. Bar graph showing the percentage of CRKP among *K. pneumoniae* genomes across the six WHO regions. B. Pie chart illustrating the contribution of each WHO region to the global CRKP isolate distribution, with genome counts indicated. Abbreviations: CRKP, carbapenem-resistant *Klebsiella pneumoniae*; WHO, World Health Organization.

**Figure S2.**
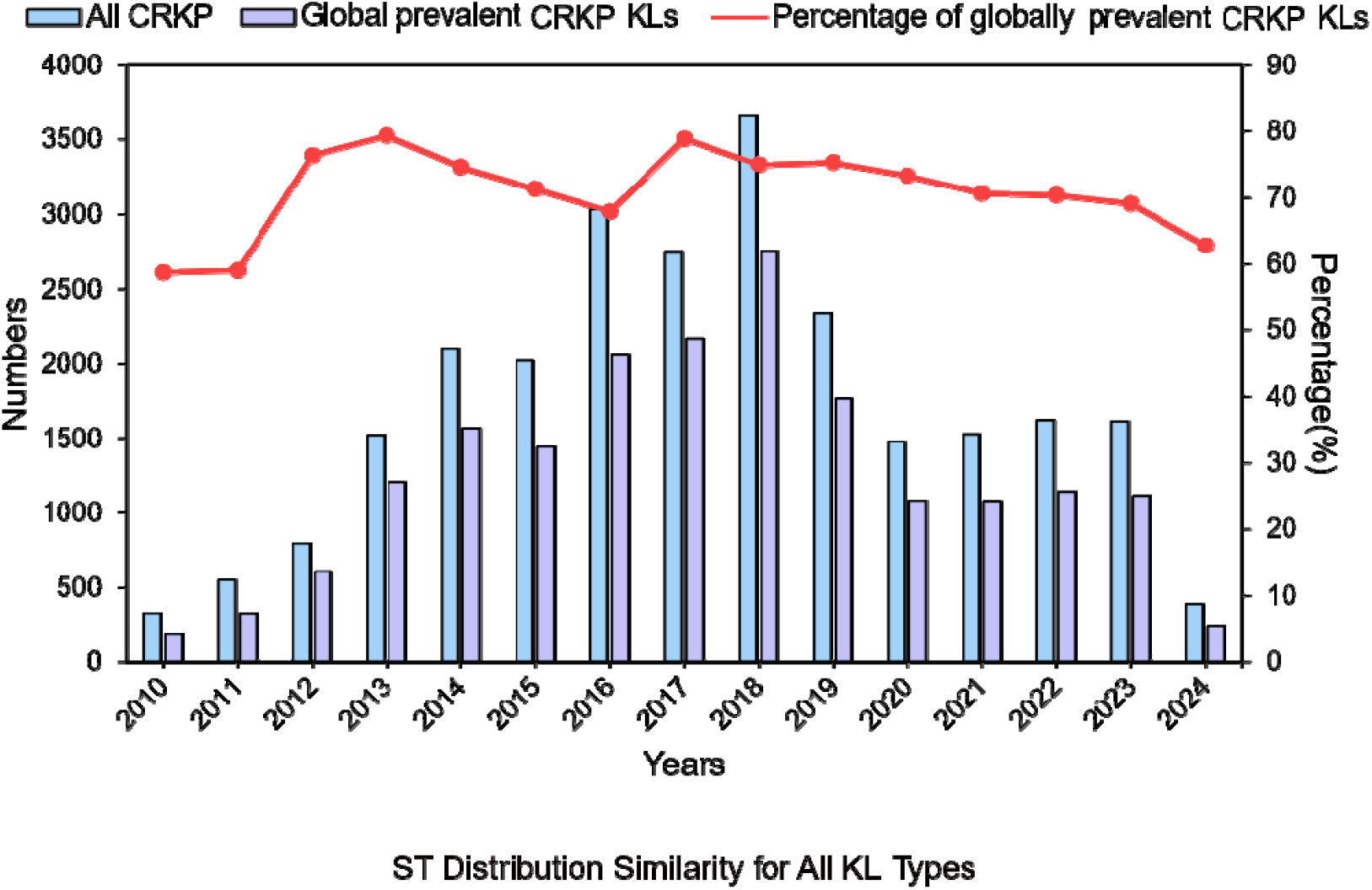
Temporal trends of globally prevalent KL types in CRKP (2010–2024). Bars (left y-axis): annual genome counts—light blue for all CRKP isolates, light purple for the globally prevalent KL subset. Line (right y-axis): yearly proportion of globally prevalent KL types within total CRKP isolates.

**Figure S3.**
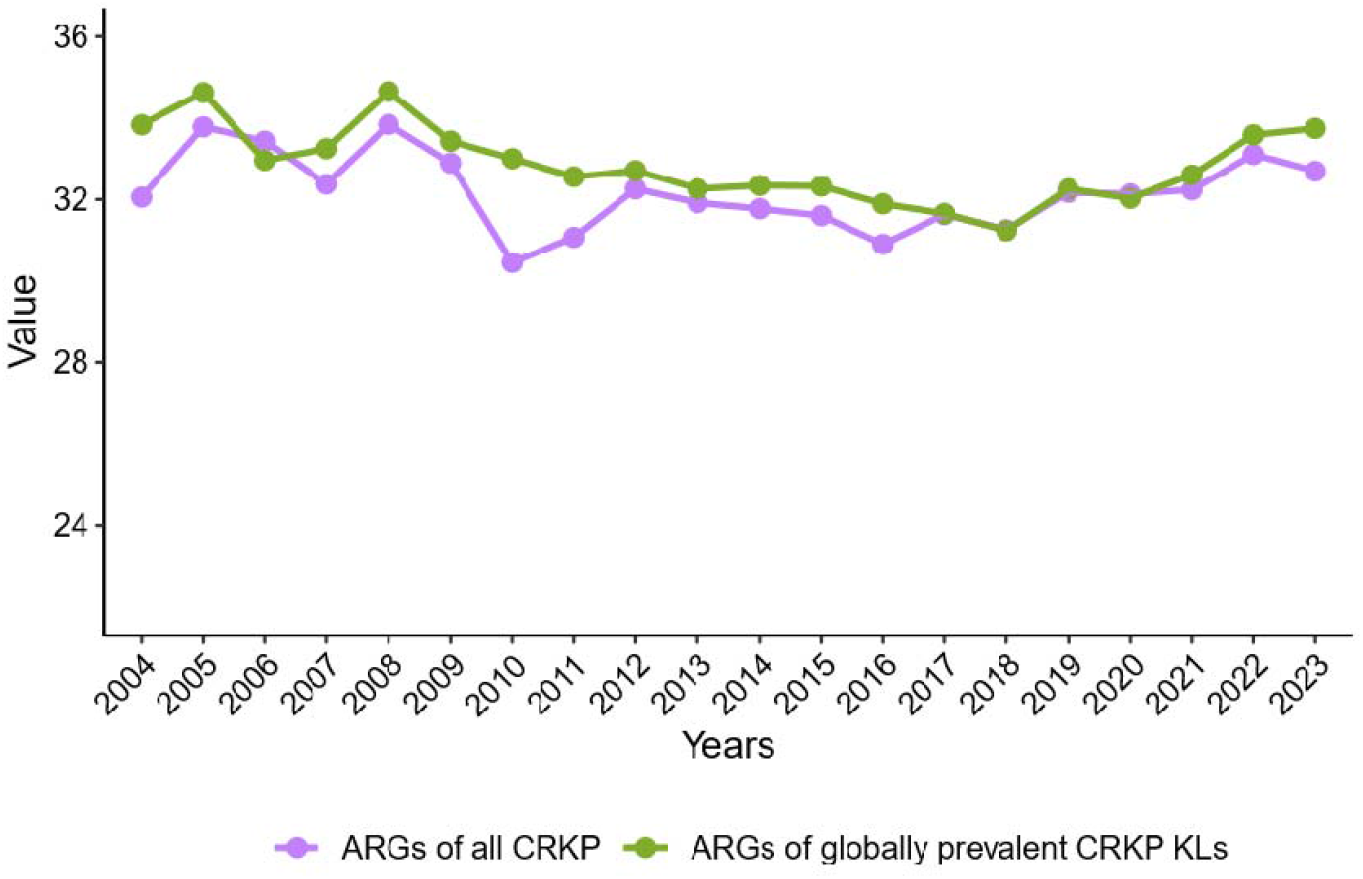
The average number of antibiotic resistance genes (ARGs) per CRKP genome from 2000 to 2023. The two series show the average number of ARGs per genome for all CRKP isolates and for the globally prevalent CRKP KL subset, respectively.

**Figure S4.**
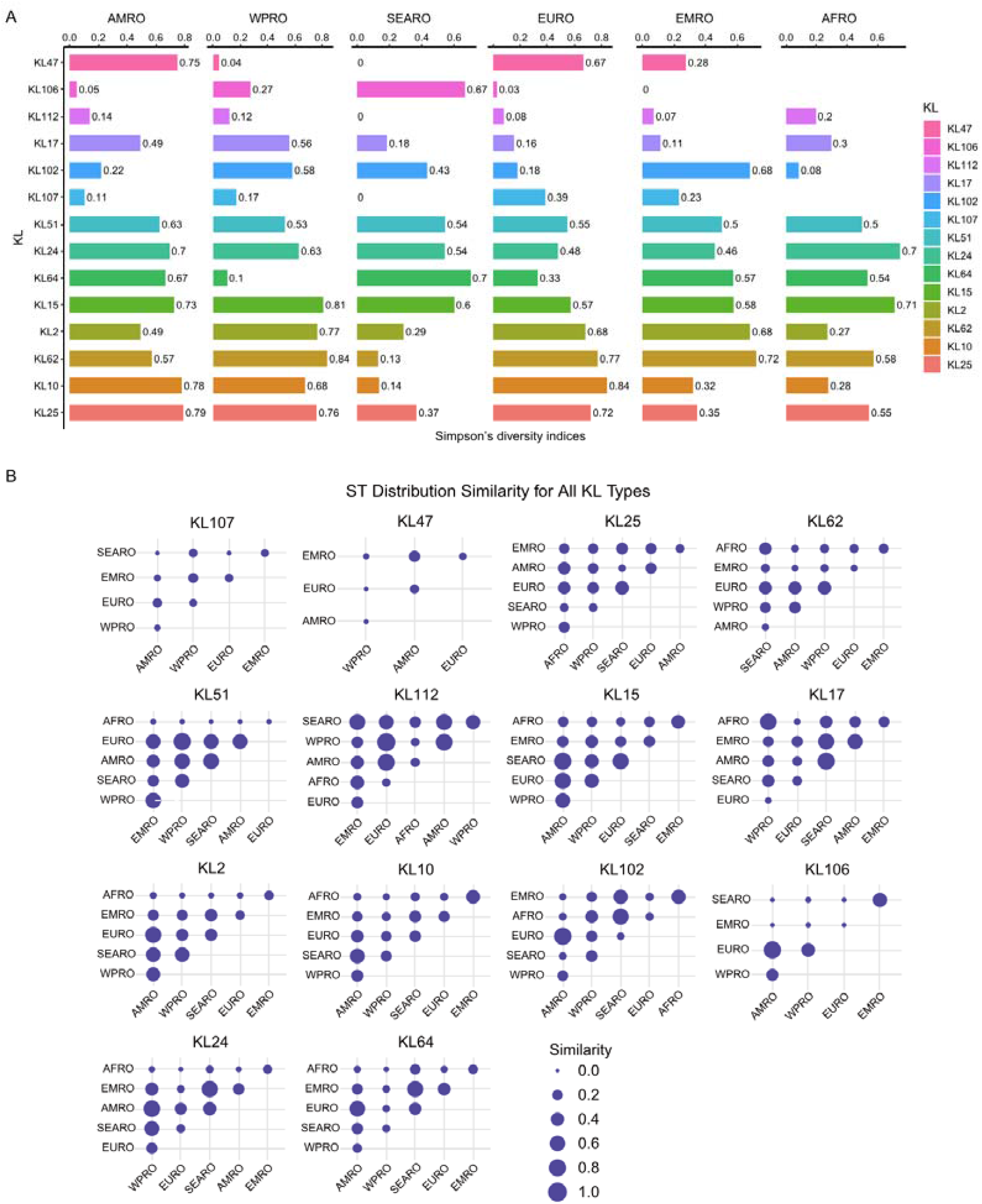
ST diversity within the six WHO regions across the 14 globally prevalent CRKP KL types. A. Simpson’s diversity indices of ST distributions in CRKP under different KL types in the six WHO regions. The 14 globally prevalent KL types are shown, with different colors representing different KL types and the length of the columns representing the size of the Simpson’s diversity indices, which are marked with numbers. B. The similarity of ST distributions among WHO regions in the 14 globally prevalent CRKP KL types. The labels on the horizontal and vertical axes represent the six WHO regions; the size of the dots indicates the degree of similarity between the ST distributions in two regions, with larger dots indicating higher similarity.

**Figure S5.**
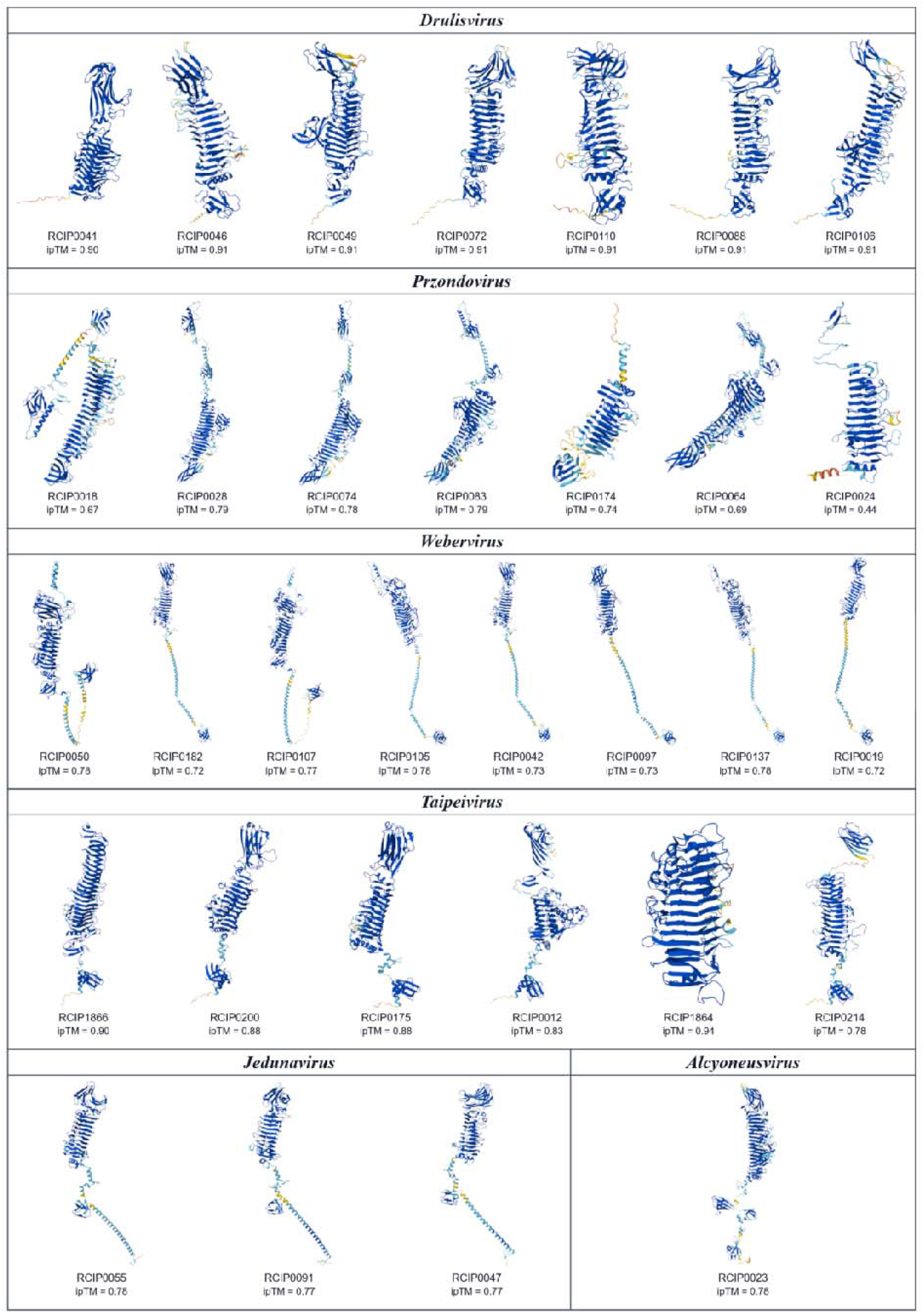
Three-dimensional structures of the predicted depolymerases. AlphaFold3-predicted monomer structures of the predicted depolymerase receptor-binding proteins (RBPs) for the 32 phages that carry a depolymerase, grouped by the six phage genera (*Drulisvirus*, *Przondovirus*, *Webervirus*, *Taipeivirus*, *Jedunavirus*, and *Alcyoneusvirus*). Each model exhibits an N-terminal anchoring domain and a central β-helical enzymatic domain, the characteristic architecture of capsular depolymerases, consistent with their function in capsular polysaccharide recognition and degradation. The interface predicted Template Modeling (ipTM) score reported by AlphaFold3 is indicated for each structure (range, 0.44–0.94).

**Figure S6.**
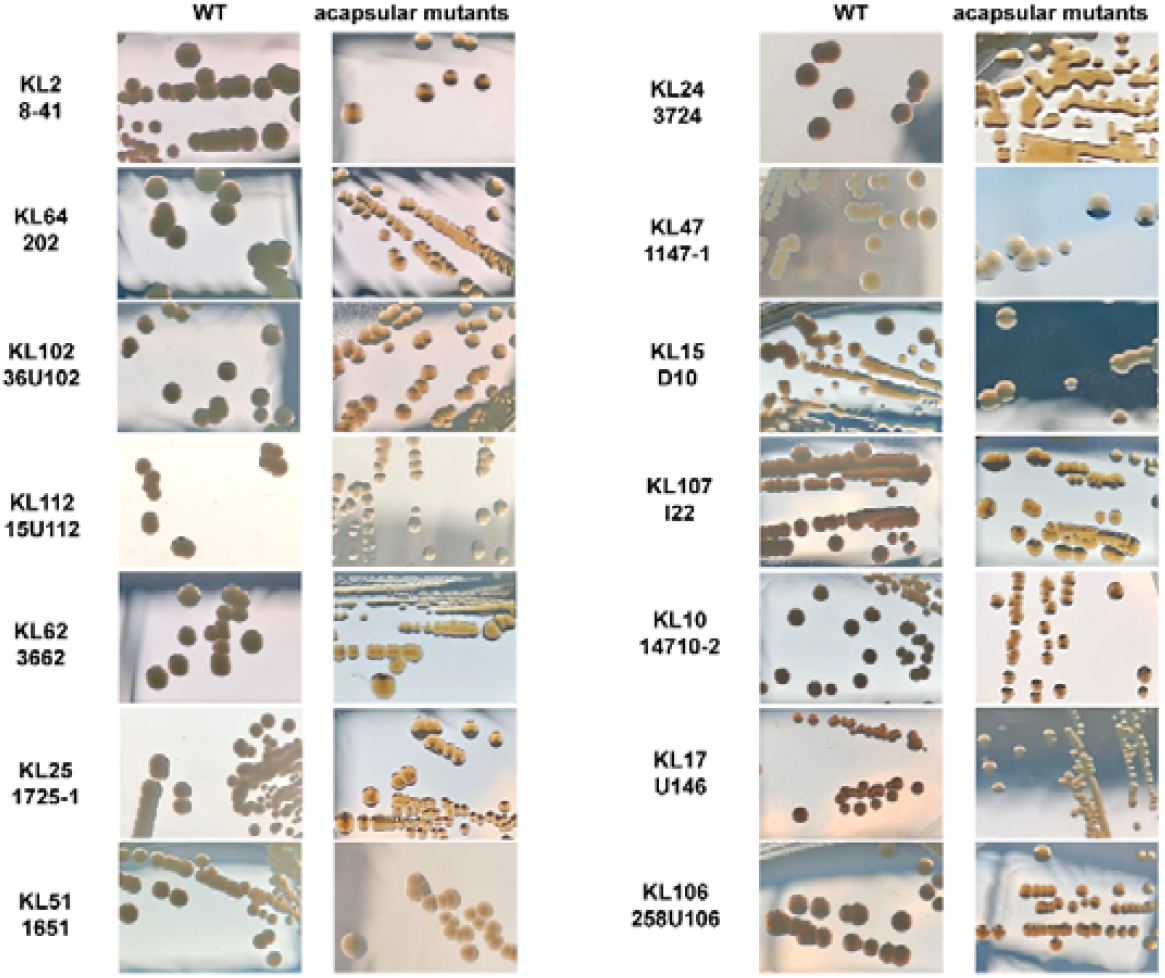
Representative colony morphology of 14 KL-type *K. pneumoniae* wild-type (WT) strains (mucoid/opaque) and their acapsular mutants (Cap, non-mucoid/translucent), indicative of capsule loss.

**Figure S7.**
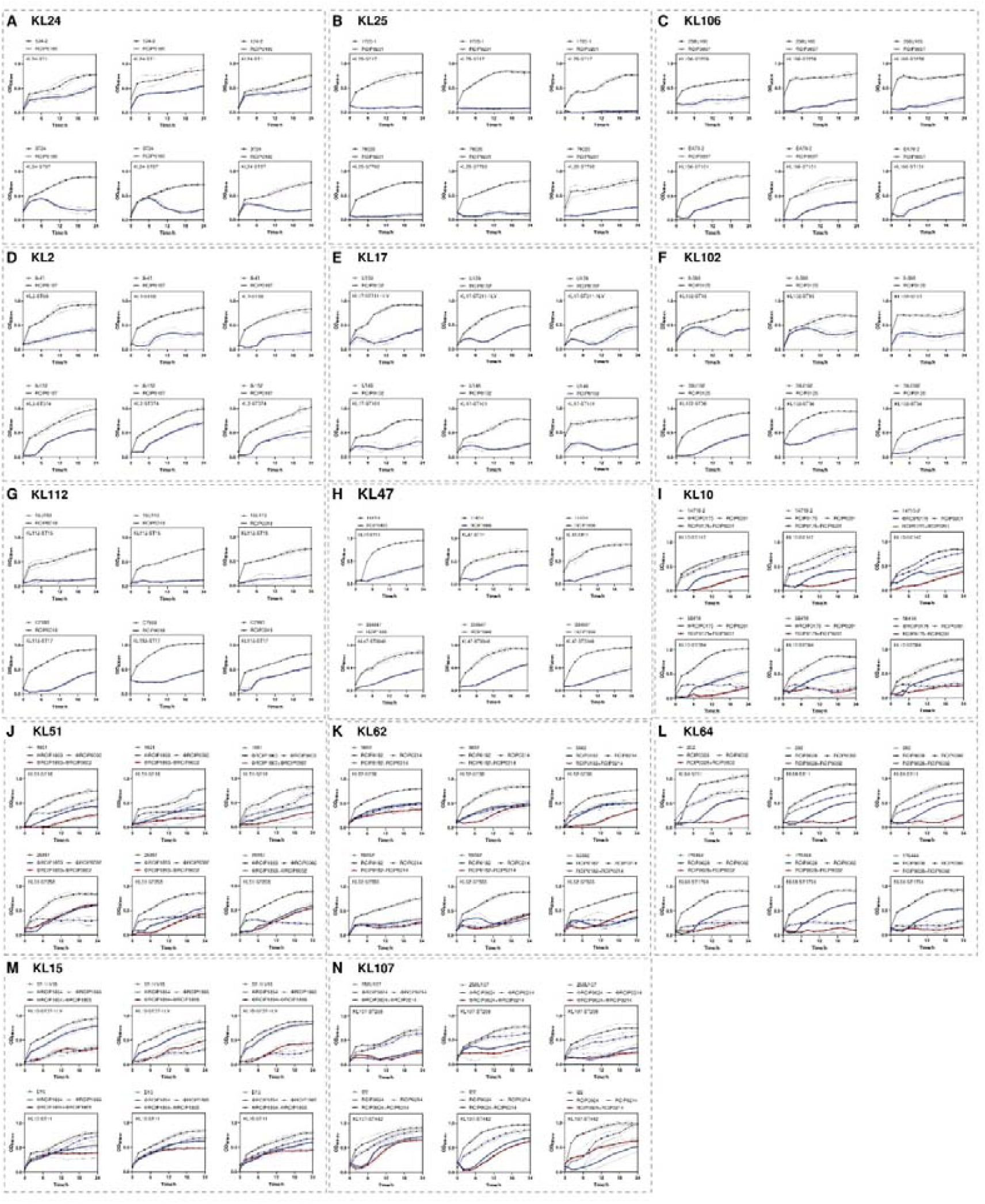
Growth curves of the 14 globally prevalent KL types represented by diverse sequence types (STs) treated with individual phages and phage cocktails in vitro. Each panel corresponds to a distinct KL type, with sub-panels showing strains of two distinct STs. Curves represent bacterial growth dynamics (OD_600_) over 24 h for the untreated control, the capsule-dependent phage (phage 1), the capsule-independent phage (phage 2), and the dual-phage cocktail (phage 1+2). Data are means ± SD of three independent replicates.

Table S1: Percentage of genomes that are potentially resistant in each KL type

Table S2: Number of ST type genomes in CRKP of various WHO regions for each globally prevalent KL

Table S3: Characteristics of the 48 sequenced *Klebsiella* phage genomes and NMDC accession numbers

Table S4: Detailed information about the 65 *K. pneumoniae* strains used for phage-bacteria infection and NMDC accession numbers

Table S5: Comparative genomic analysis of acapsular (Cap) mutants derived from 14 KL types, revealing mutations associated with capsule biosynthesis.

Table S6: Comparative genomic analysis of KL64 mutants derived from a two-step strain with defects in CPS and LPS.

## Notes

### Competing Interest Statement

The authors have declared no competing interest.

