## Supplementary material for "Capsular surveillance-guided phage cocktails against carbapenem-resistant *Klebsiella pneumoniae*": FigS1

A

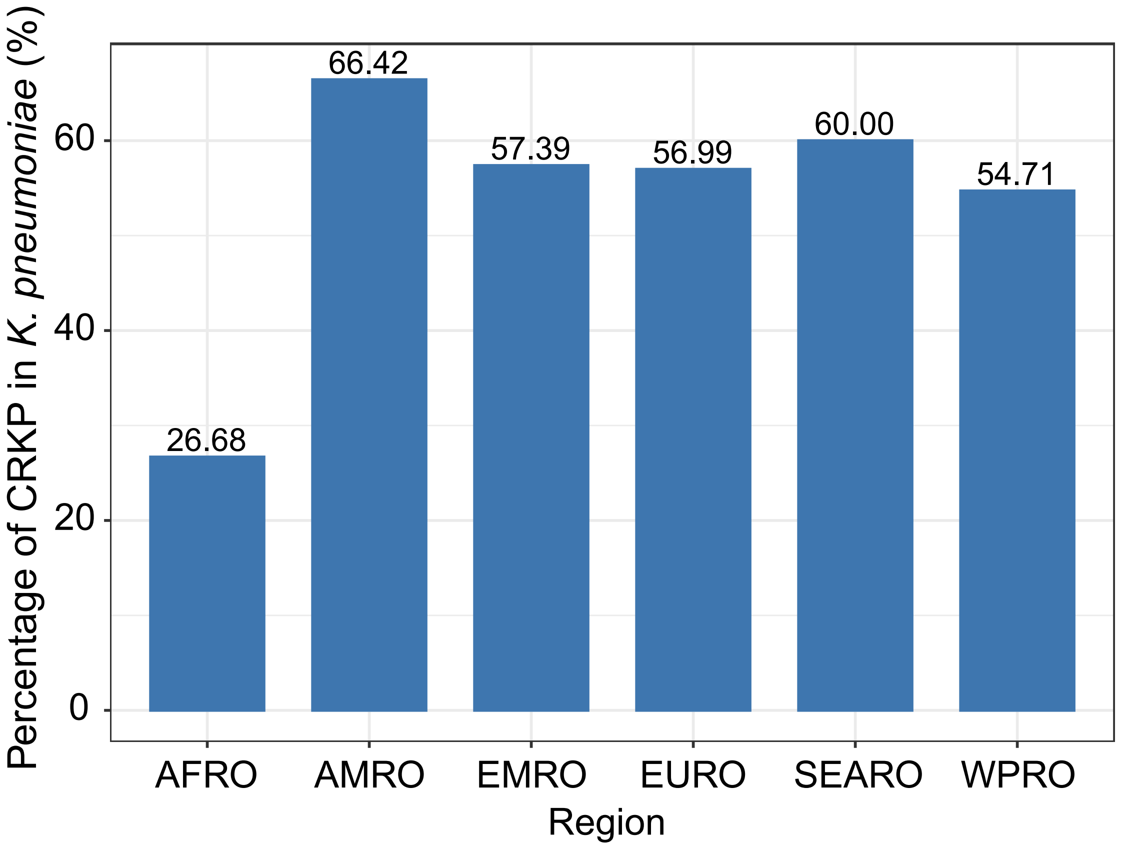

Percentage of CRKP in K. pneumoniae (%)
0
20
40
60
26.68
66.42
57.39
56.99
60.00
54.71
AFRO
AMRO
EMRO
EURO
SEARO
WPRO
Region


B

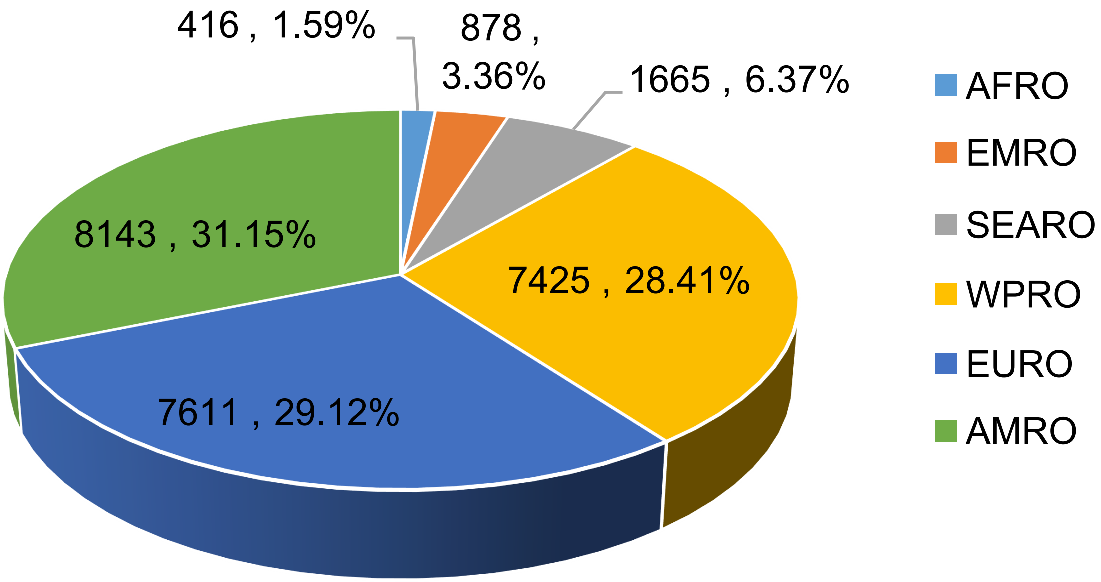

416 , 1.59%
878 , 3.36%
1665 , 6.37%
8143 , 31.15%
7425 , 28.41%
7611 , 29.12%
AFRO
EMRO
SEARO
WPRO
EURO
AMRO
