## Supplementary figures and images for "Capsular surveillance-guided phage cocktails against carbapenem-resistant *Klebsiella pneumoniae*"

### FigS2

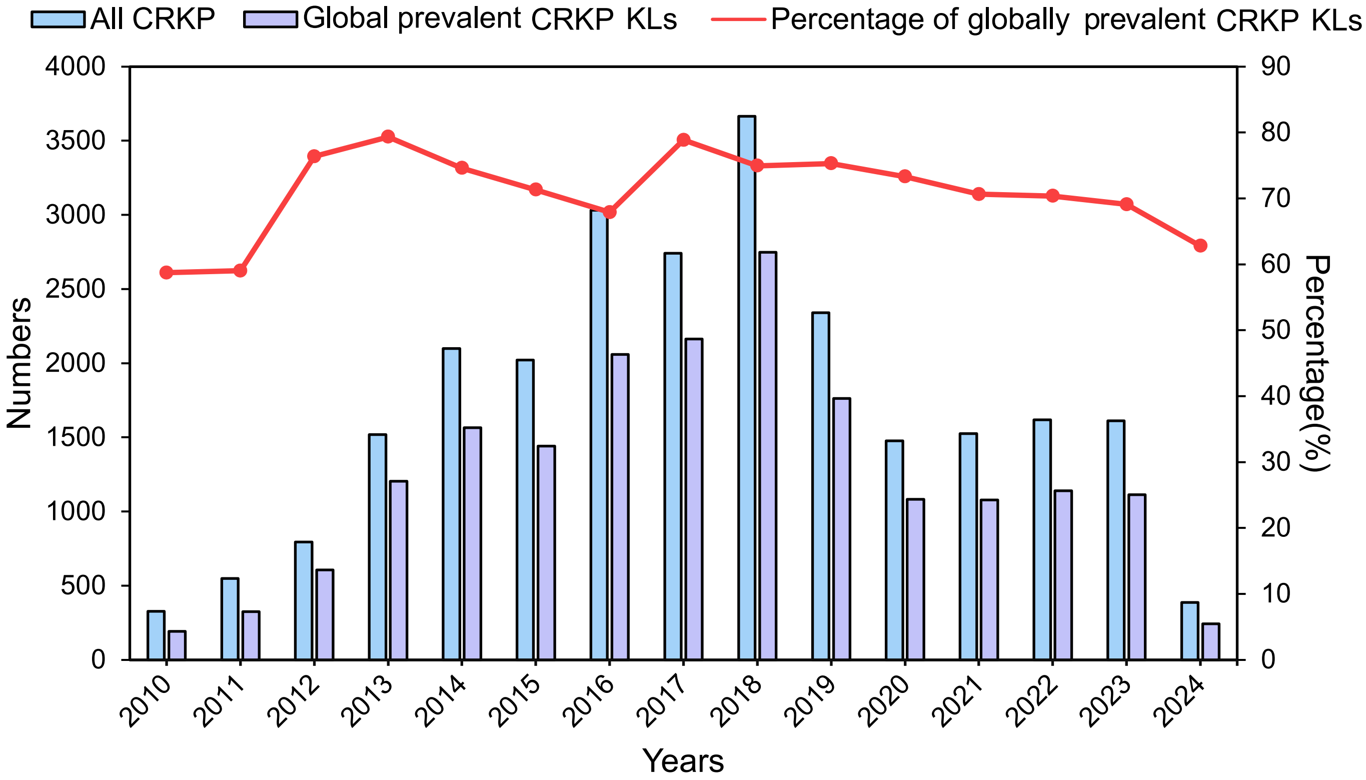


ST Distribution Similarity for All KL Types

### FigS3

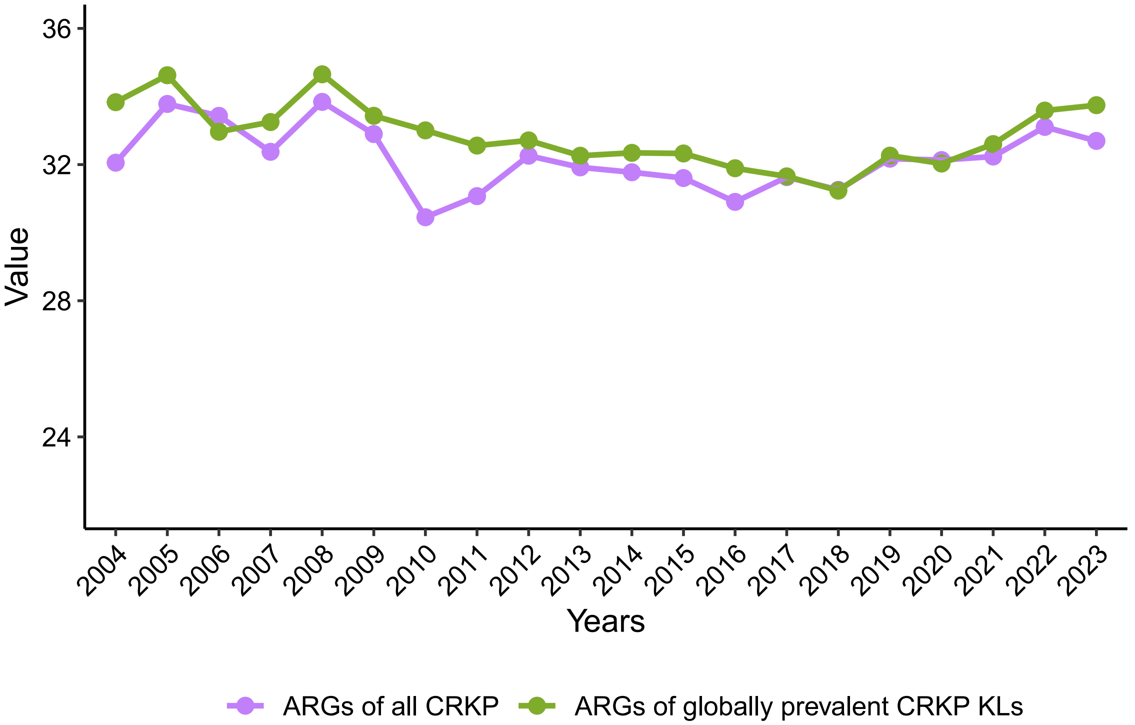

Value
36
32
28
24
2004
2005
2006
2007
2008
2009
2010
2011
2012
2013
2014
2015
2016
2017
2018
2019
2020
2021
2022
2023
Years
ARGs of all CRKP
ARGs of globally prevalent CRKP KLs

### FigS6

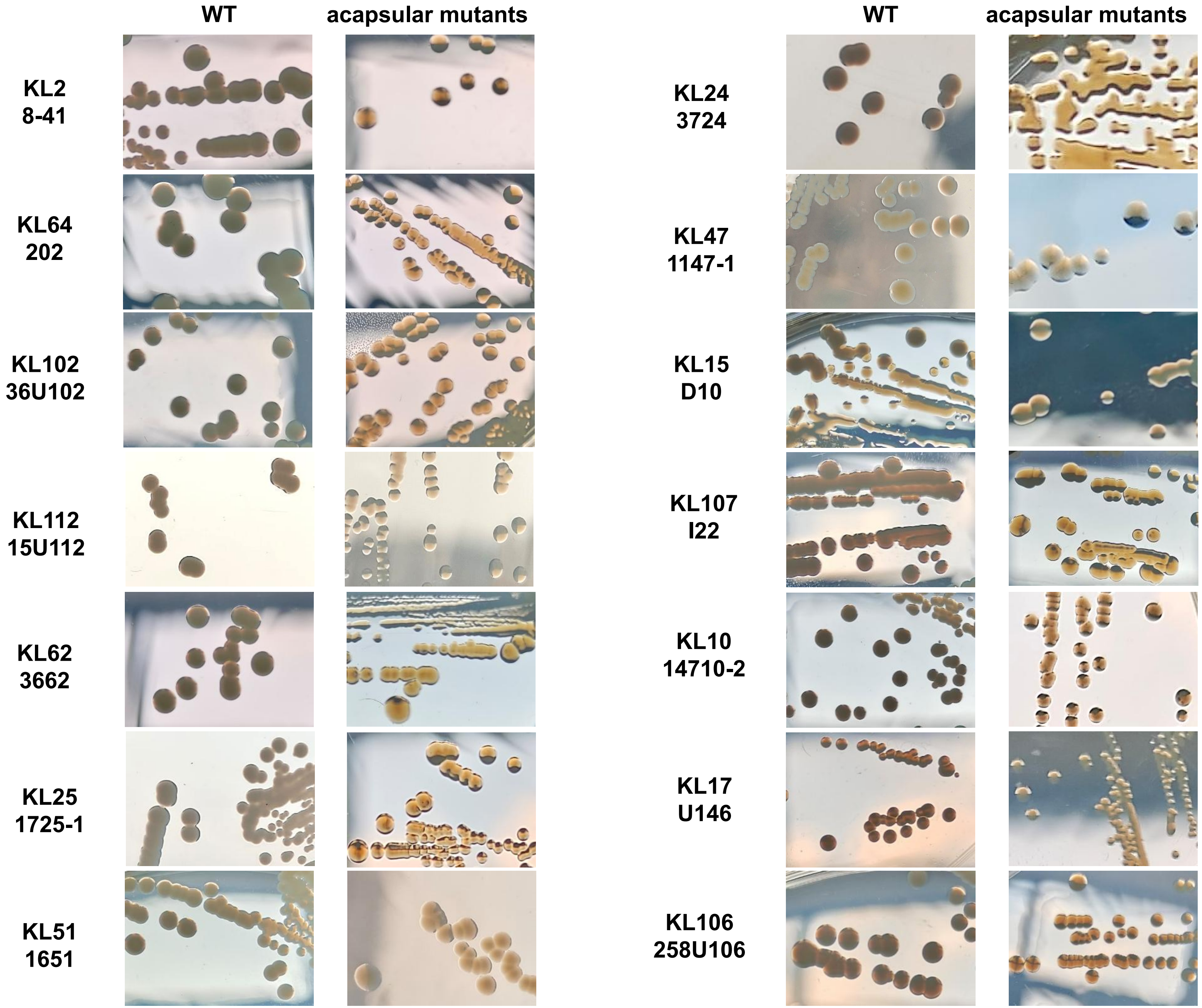
WT
acapsular mutants
KL2
8-41
KL64
202
KL102
36U102
KL112
15U112
KL62
3662
KL25
1725-1
KL51
1651
WT
acapsular mutants
KL24
3724
KL47
1147-1
KL15
D10
KL107
I22
KL10
14710-2
KL17
U146
KL106
258U106
