## Supplementary material for "Capsular surveillance-guided phage cocktails against carbapenem-resistant *Klebsiella pneumoniae*": FigS4

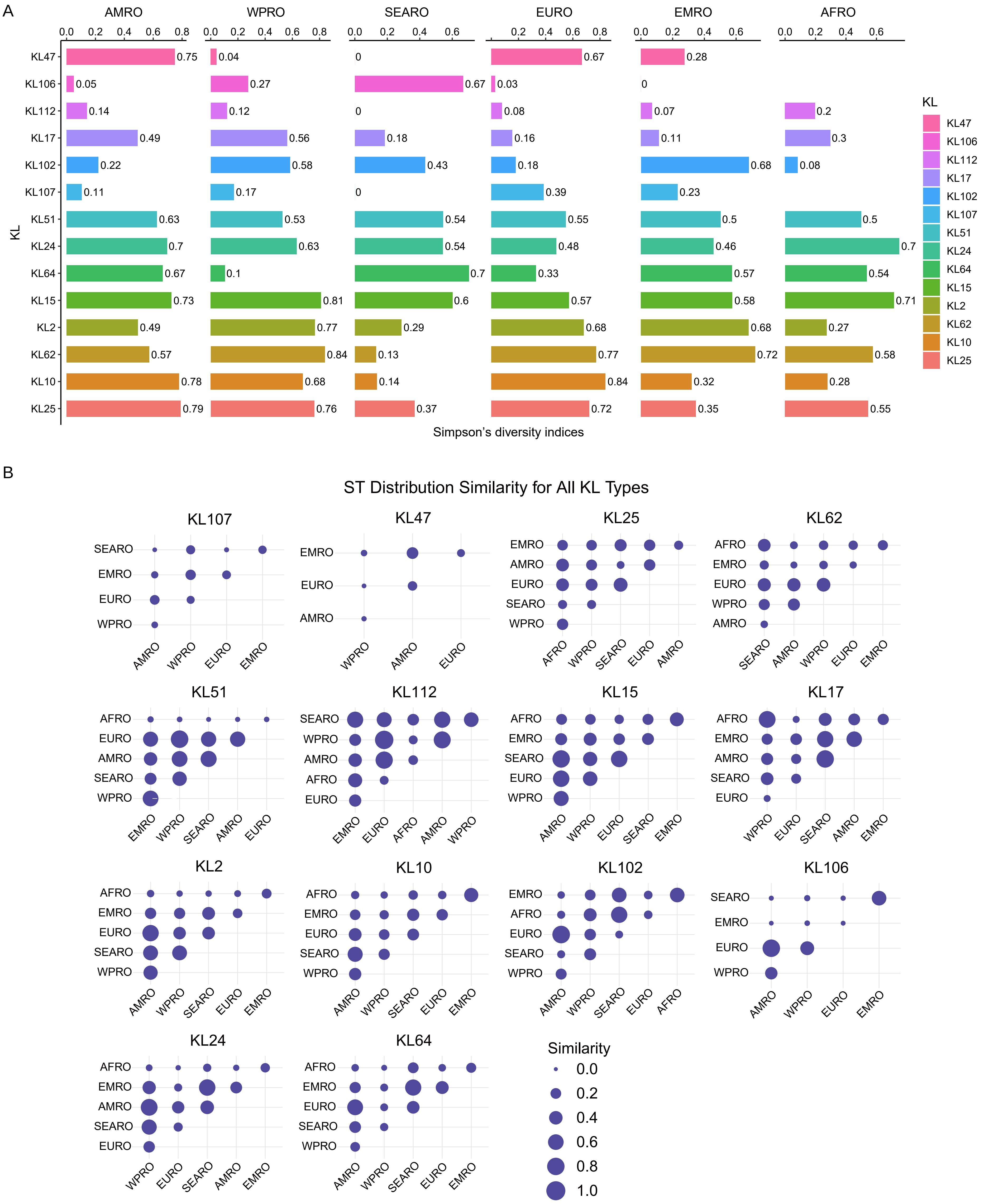
A
AMRO
WPRO
SEARO
EURO
EMRO
AFRO
KL
KL47
KL106
KL112
KL17
KL102
KL107
KL51
KL24
KL64
KL15
KL2
KL62
KL10
KL25
AMRO: 0.75 0.05 0.14 0.49 0.22 0.11 0.63 0.7 0.67 0.73 0.49 0.57 0.78 0.79
WPRO: 0.04 0.27 0.12 0.56 0.58 0.17 0.53 0.63 0.1 0.81 0.77 0.84 0.68 0.76
SEARO: 0 0.67 0 0.18 0.43 0 0.54 0.54 0.7 0.6 0.29 0.13 0.14 0.37
EURO: 0.67 0.03 0.08 0.16 0.18 0.39 0.55 0.48 0.33 0.57 0.68 0.77 0.84 0.72
EMRO: 0.28 0 0.07 0.11 0.68 0.23 0.5 0.46 0.57 0.58 0.68 0.72 0.32 0.35
AFRO: 0.2 0.3 0.08 0.5 0.7 0.54 0.71 0.27 0.58 0.28 0.55
Simpson's diversity indices
B
ST Distribution Similarity for All KL Types
KL107
KL47
KL25
KL62
KL51
KL112
KL15
KL17
KL2
KL10
KL102
KL106
KL24
KL64
AMRO
WPRO
SEARO
EURO
EMRO
AFRO
Similarity
0.0
0.2
0.4
0.6
0.8
1.0
