## Supplementary material for "Capsular surveillance-guided phage cocktails against carbapenem-resistant *Klebsiella pneumoniae*": FigS5

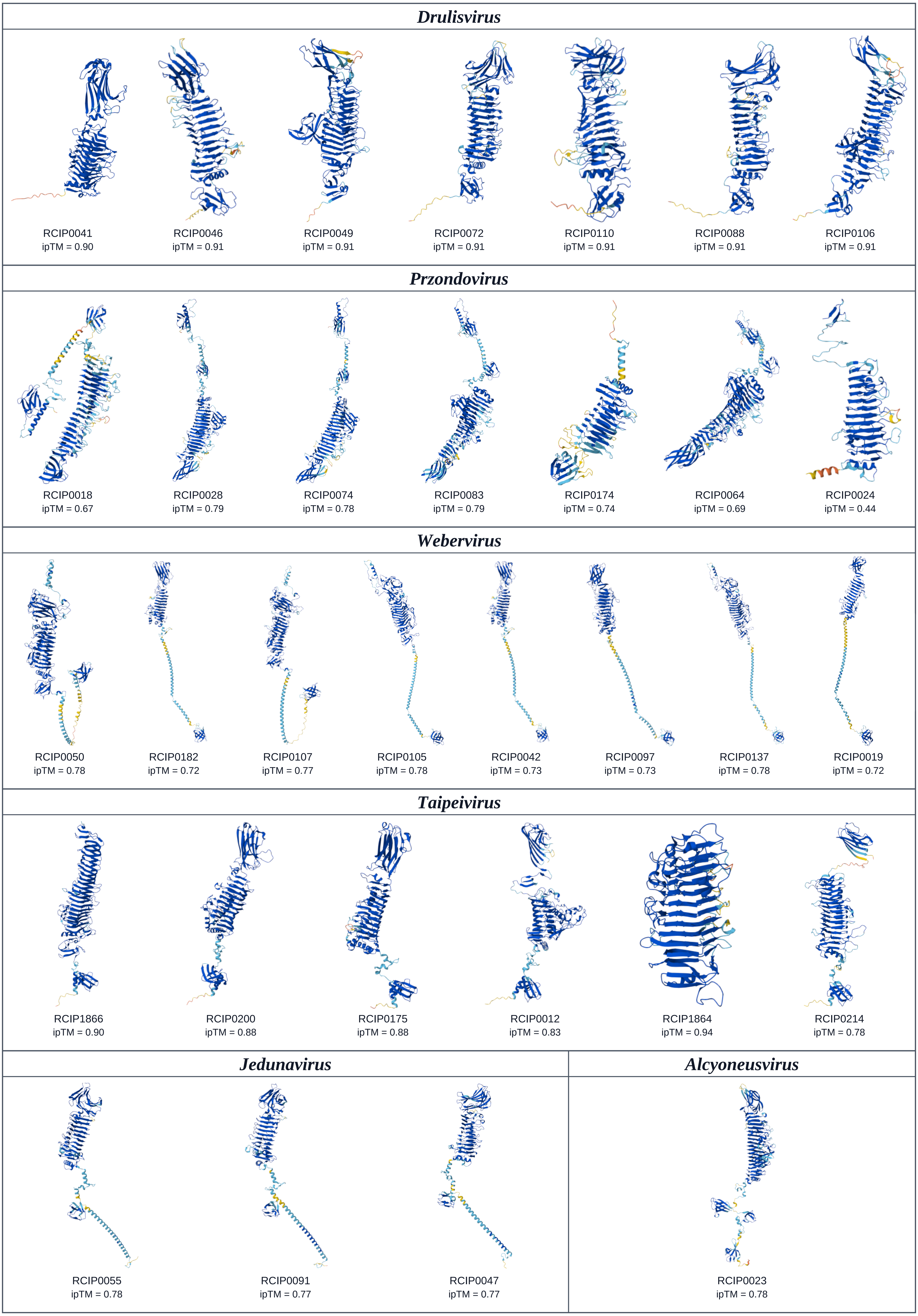
Drulisvirus
RCIP0041
ipTM = 0.90
RCIP0046
ipTM = 0.91
RCIP0049
ipTM = 0.91
RCIP0072
ipTM = 0.91
RCIP0110
ipTM = 0.91
RCIP0088
ipTM = 0.91
RCIP0106
ipTM = 0.91
Przondovirus
RCIP0018
ipTM = 0.67
RCIP0028
ipTM = 0.79
RCIP0074
ipTM = 0.78
RCIP0083
ipTM = 0.79
RCIP0174
ipTM = 0.74
RCIP0064
ipTM = 0.69
RCIP0024
ipTM = 0.44
Webervirus
RCIP0050
ipTM = 0.78
RCIP0182
ipTM = 0.72
RCIP0107
ipTM = 0.77
RCIP0105
ipTM = 0.78
RCIP0042
ipTM = 0.73
RCIP0097
ipTM = 0.73
RCIP0137
ipTM = 0.78
RCIP0019
ipTM = 0.72
Taipeivirus
RCIP1866
ipTM = 0.90
RCIP0200
ipTM = 0.88
RCIP0175
ipTM = 0.88
RCIP0012
ipTM = 0.83
RCIP1864
ipTM = 0.94
RCIP0214
ipTM = 0.78
Jedunavirus
RCIP0055
ipTM = 0.78
RCIP0091
ipTM = 0.77
RCIP0047
ipTM = 0.77
Alcyoneusvirus
RCIP0023
ipTM = 0.78
