## Supplementary material for "Capsular surveillance-guided phage cocktails against carbapenem-resistant *Klebsiella pneumoniae*": FigS7

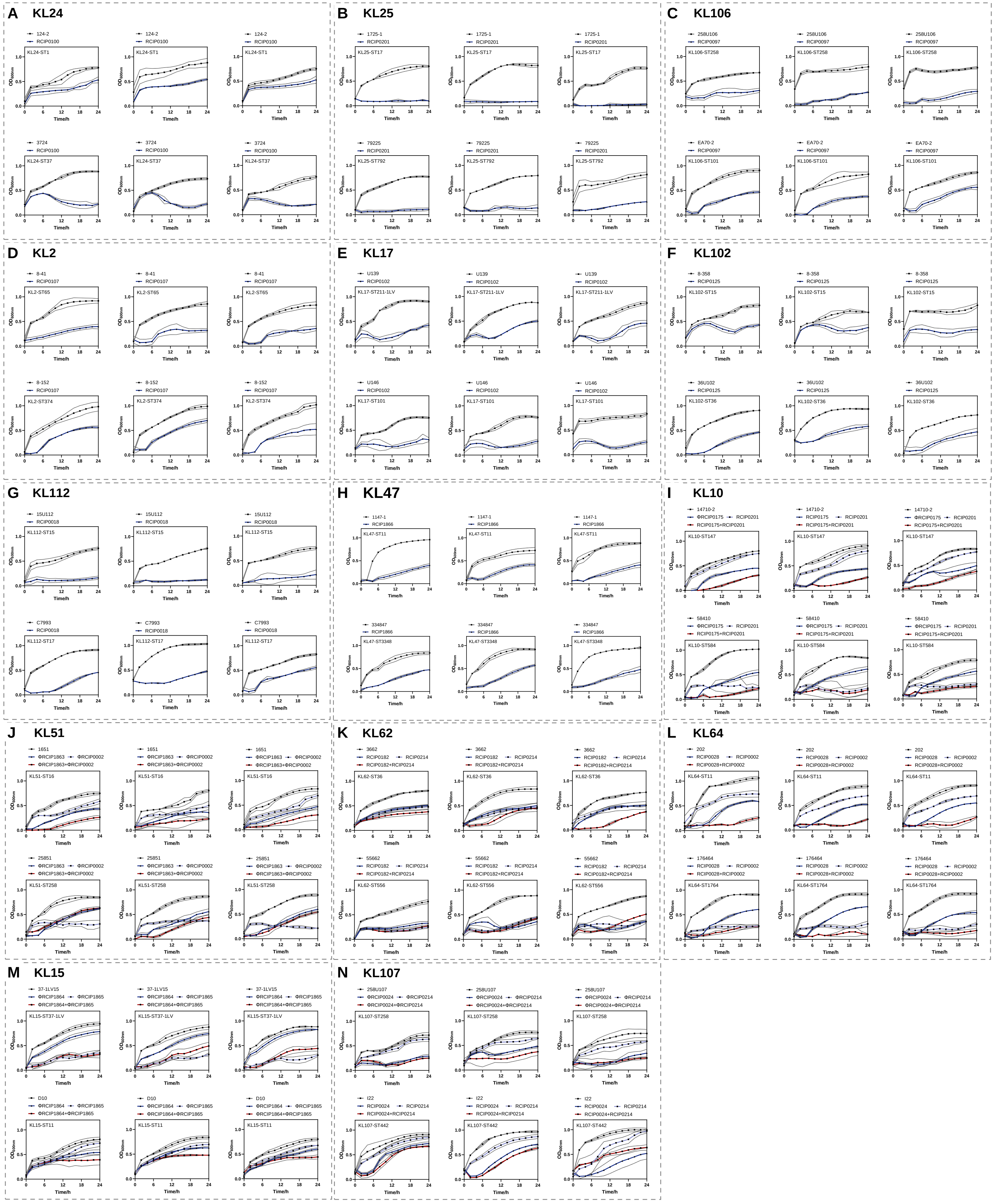

A KL24
124-2
RCIP0100
KL24-ST1
3724
KL24-ST37
B KL25
1725-1
RCIP0201
KL25-ST17
79225
KL25-ST792
C KL106
258U106
RCIP0097
KL106-ST258
EA70-2
KL106-ST101
D KL2
8-41
RCIP0107
KL2-ST65
8-152
KL2-ST374
E KL17
U139
RCIP0102
KL17-ST211-1LV
U146
KL17-ST101
F KL102
8-358
RCIP0125
KL102-ST15
36U102
KL102-ST36
G KL112
15U112
RCIP0018
KL112-ST15
C7993
KL112-ST17
H KL47
1147-1
RCIP1866
KL47-ST11
334847
KL47-ST3348
I KL10
14710-2
ΦRCIP0175
RCIP0201
RCIP0175+RCIP0201
KL10-ST147
58410
KL10-ST584
J KL51
1651
ΦRCIP1863
ΦRCIP0002
ΦRCIP1863+ΦRCIP0002
KL51-ST16
25851
KL51-ST258
K KL62
3662
RCIP0182
RCIP0214
RCIP0182+RCIP0214
KL62-ST36
55662
KL62-ST556
L KL64
202
RCIP0028
RCIP0002
RCIP0028+RCIP0002
KL64-ST11
176464
KL64-ST1764
M KL15
37-1LV15
ΦRCIP1864
ΦRCIP1865
ΦRCIP1864+ΦRCIP1865
KL15-ST37-1LV
D10
KL15-ST11
N KL107
258U107
ΦRCIP0024
ΦRCIP0214
ΦRCIP0024+ΦRCIP0214
KL107-ST258
I22
RCIP0024
RCIP0214
RCIP0024+RCIP0214
KL107-ST442
OD600nm
Time/h
